# A systematic approach identifies p53-DREAM target genes associated with blood or brain abnormalities

**DOI:** 10.1101/2023.06.21.545932

**Authors:** Jeanne Rakotopare, Vincent Lejour, Carla Duval, Eliana Eldawra, Hugues Escoffier, Franck Toledo

## Abstract

p53 is mainly known as a tumor suppressor, but mouse models revealed that increased p53 activity may cause bone marrow failure, through mechanisms that likely include gene repression mediated by the p53-DREAM pathway. Here we designed a systematic approach to identify p53-DREAM targets whose repression might contribute to abnormal hematopoiesis. We used gene ontology to analyze transcriptomic changes associated with bone marrow cell differentiation and p53 activation, then ChIP-seq data to find promoters bound by the DREAM complex. We next created positional frequency matrices to identify evolutionary conserved sequence elements potentially bound by DREAM. The same approach was developed to find p53-DREAM targets associated with brain abnormalities, also observed in mice with increased p53 activity. Putative DREAM binding sites were found for 151 candidate p53-DREAM target genes, of which 106 are mutated in a blood or brain genetic disorder. Twenty-one DREAM binding sites were tested and found to impact on gene expression in luciferase reporter assays, notably regulating genes mutated in dyskeratosis congenita (*Rtel1*), Fanconi anemia (*Fanca*), Diamond-Blackfan anemia (*Tsr2*), primary microcephaly (*Casc5*, *Ncaph*, *Wdr62*) or pontocerebellar hypoplasia (*Toe1*). These results provide clues on the role of the p53-DREAM pathway in regulating hematopoiesis and brain development, with implications for tumorigenesis.

**Author Summary:** The capacity of p53 to activate the transcription of genes important for cell cycle arrest, apoptosis or cellular metabolism has been recognized for decades. By contrast, the potential importance of p53-dependent transcriptional repression emerged more recently. Although p53 frequently appears to repress genes indirectly via the DREAM repressor complex, only a few studies attempted to define p53-DREAM target gene repertoires, often by analyzing cell cycle regulation in fibroblasts. Here we aimed to gain a better appreciation of the clinical relevance of the p53-DREAM pathway by designing a systematic approach for the identification of p53-DREAM targets. Because mouse models with increased p53 activity suffer from bone marrow failure or brain hypoplasia, we relied on transcriptomic changes associated with bone marrow cell differentiation, blood- and brain-related gene ontology terms and RNAseq data from hematopoietic or neural progenitor cells to identify p53-DREAM targets, then created positional frequency matrices to find putative DREAM binding sites. Our study provides a resource of predicted DREAM binding sites for 151 genes associated with blood and/or brain abnormalities, many of which were not previously known to be DREAM targets. Furthermore, our analysis suggests that p53-DREAM alterations may contribute to phenotypic variations in glioblastoma cells.

## Introduction

The <u>d</u>imerization partner, <u>R</u>B-like, <u>E</u>2F4/5 <u>a</u>nd <u>M</u>uvB (DREAM) complex is a master coordinator of cell-cycle dependent gene expression, that mediates gene repression in quiescent cells [1] and coordinates periodic gene expression in proliferating cells [2]. Although p53 was shown to repress transcription over 30 years ago [3,4], its capacity to do so indirectly, via p21 and DREAM, emerged only progressively [5–11]. Meta-analyses first indicated that the p53-p21-DREAM pathway regulates G2/M cell cycle genes [12], then that it participates in the control of all cell cycle checkpoints [13,14], and 85% of known targets of the p53-p21-RB pathway were recently proposed to be also regulated by p53-p21-DREAM signaling [15]. Furthermore cells lacking LIN37, a subunit of the DREAM complex, demonstrated the functional impact of the p53-p21-DREAM (called below p53-DREAM) pathway in cell cycle regulation [16,17].

However, the relative importance of this pathway remains to be fully appreciated, because multiple mechanisms were proposed to account for p53-mediated gene repression [18]. In fact, hundreds of genes were proposed to be regulated by the p53-DREAM pathway, but so far only a few DREAM binding sites were demonstrated experimentally, perhaps due to the complexity of DREAM binding. The DREAM complex was initially reported to repress the transcription of genes whose promoter sequences contain a bipartite binding motif called CDE/CHR [19,20] (or E2F/CHR [21]), with a GC-rich <u>c</u>ell cycle <u>d</u>ependent <u>e</u>lement (CDE) that may be bound by E2F4 or E2F5, and an AT-rich <u>c</u>ell cycle gene <u>h</u>omology <u>r</u>egion (CHR) that may be bound by LIN54, the DNA-binding subunit of MuvB [19,20]. Later studies indicated that DREAM may also bind promoters with a single E2F binding site, a single CHR element, or a bipartite E2F/<u>C</u>HR-like <u>e</u>lement (CLE), and concluded that E2F and CHR elements are required for the regulation of G1/S and G2/M cell cycle genes, respectively [14,22]. The Target gene regulation database of p53 and cell-cycle genes [23] was reported to include putative DREAM binding sites for many human genes, based on separate genome-wide searches for 7 bp-long E2F or 5 bp-long CHR motifs, but the predicted sites were not tested experimentally. By contrast, positional frequency matrices designed to find bipartite DREAM binding sites were used to analyze only a few promoters, but their predictions were confirmed experimentally [24,25].

Our interest in the p53-DREAM pathway stems from the analysis of a mouse model with increased p53 activity. We observed that mutant mice expressing p53^Δ31^, a truncated protein lacking 31 residues of the C-terminal domain, exhibited all the phenotypic traits associated with dyskeratosis congenita and its severe variant the Hoyeraal-Hreidarsson syndrome, two bone marrow failure syndromes caused by defective telomere maintenance [26]. Accordingly, p53^Δ31/Δ31^ mice exhibited short telomeres and a reduced expression of a few genes mutated in dyskeratosis congenita, notably *Rtel1*, whose expression levels correlated with mouse survival [26]. p53^Δ31/Δ31^ cells also exhibited a reduced capacity to repair DNA interstrand crosslinks, a typical feature of cells from patients with Fanconi anemia, another bone marrow failure syndrome [25]. This phenotype could be explained by a reduced expression of several genes of the Fanconi anemia DNA repair pathway, including the *Fancd2*, *Fanci* and *Rad51* genes, whose promoters contain functionally relevant bipartite DREAM binding sites [25]. These findings appeared potentially relevant to human pathological processes, because p53 could also repress the homologous human genes [25,26]. In agreement with this, we later identified a germline missense mutation of *MDM4*, encoding a major p53 negative regulator, in a familial syndrome of neutropenia and defective telomere maintenance, and could correlate p53 activation with decreased *RTEL1* expression and short telomeres in the most affected family member, as well as in mice carrying the same *Mdm4* mutation [27]. Furthermore, two individuals carrying germline *TP53* mutations leading to express a truncated p53 protein lacking 32 C-terminal residues were recently reported [28]. Consistent with our findings, these individuals exhibited increased p53 activity and short telomeres. Interestingly however, they suffered from a pure red cell aplasia resembling Diamond-Blackfan anemia, another bone marrow failure syndrome caused by ribosomal dysfunction - although the molecular mechanisms underlying impaired erythrocyte production in these patients remained unexplained [28]. Taken together, these data indicated that germline p53 activation can cause a large spectrum of phenotypic traits found in patients with either dyskeratosis congenita, Fanconi anemia or Diamond-Blackfan anemia.

Our results in p53^Δ31/Δ31^ mice led us to hypothesize that these phenotypic traits might result from gene repression mediated by the p53-DREAM pathway, which incited us to design a genome-wide approach relying on gene ontology and bone marrow cell differentiation to identify p53-DREAM targets related to hematopoiesis. Furthermore, mice and humans with germline increases in p53 activity may also exhibit microcephaly or cerebellar hypoplasia [26,28], and cerebellar hypoplasia may be observed in a subset of patients with bone marrow failure syndromes, including patients with the Hoyeraal-Hreidarsson syndrome [29,30] or 27% of patients with Fanconi anemia [31]. This led us to use the same strategy to search for candidate p53-DREAM target genes that might be involved in brain abnormalities.

With this study we aimed to gain a better appreciation of the clinical relevance of the p53-DREAM pathway. We developed refined positional frequency matrices and identified bipartite DREAM binding sites in the promoters of 151 genes, many of which were not previously known to be DREAM targets. Most putative DREAM binding sites mapped at the level of ChIP peaks for DREAM subunits and near transcription start sites, and a subset of the sites were tested with luciferase assays. Our study provides a resource of predicted DREAM binding sites for genes associated with blood and brain abnormalities, as well as a method that might be applied to analyze genes associated with other pathologies.

## Results

### Candidate p53-DREAM target genes associated with blood abnormalities

To investigate the role of the p53-DREAM pathway on the regulation of hematopoiesis, we exploited a transcriptomic approach in bone marrow cells (BMCs). The Homeobox (Hox) family of transcription factors controls the proliferation, differentiation and self-renewal of hematopoietic stem cells. Notably, Hoxa9 is required for myeloid, erythroid and lymphoid hematopoiesis [32] and its overexpression causes hematopoietic stem cell expansion [33]. Muntean *et al.* generated a cellular model for Hoxa9 conditional expression [34]. In this model, murine bone marrow stem and progenitor cells were immortalized by transduction with Hoxa9-ER in the presence of tamoxifen, and tamoxifen withdrawal led to their differentiation within 5 days. We observed that p53 activation correlated with cell differentiation in this system, because genes known to be transactivated by p53 (e.g. *Cdkn1a*, *Mdm2*) were induced, whereas genes repressed by p53 (e.g. *Rtel1*, *Fancd2*) were downregulated after tamoxifen withdrawal (Figure 1a; see also Figure S1 for additional examples of p53-regulated genes). Thus, to investigate the impact of p53 on telomere biology we performed a gene ontology (GO) analysis of the expression data obtained with this system (Gene Expression Omnibus #GSE21299), which relied on 45,101 microarray probes corresponding to 20,627 genes, of which 17,461 are associated with a GO term according to the Gene Ontology enRIchment anaLysis and visuaLizAtion tool (GOrilla) [35]. We focused on genes downregulated at least 1.5-fold upon tamoxifen withdrawal. Such a downregulation was observed for 6,880 probes, corresponding to 3,631 genes associated with a GO term. According to the GOrilla tool, a significant enrichment was observed for 13 GO terms related to telomere biology (Table 1). These 13 GO terms are partially overlapping and correspond to 68 different genes, including 6 genes (*Brca2, Dkc1*, *Gar1*, *Rad51, Rtel1*, *Terf1*) we previously reported to be downregulated by p53 [25,26]. In addition, among the genes downregulated upon BMC differentiation, we noticed two genes (*Tyms, Zcchc8*) recently associated with genetic disorders of telomere biology [36,37], two genes (*Shq1*, *Son*) that may also impact on telomere maintenance and four p53-regulated genes (*Dek*, *Fancd2*, *Fen1*, *Timeless*) included in DNA repair GO terms but that also impact on telomeres [25]. In sum, BMC differentiation correlated with the decreased expression of 76 genes that may impact on telomere biology (Figure 1b, Table S1). Consistent with the notion that BMC differentiation strongly correlates with p53 activation in this system, 72 of these 76 genes have negative p53 expression score(s) in the Target gene regulation (TGR) database [23], which indicates that they were downregulated upon p53 activation in most experiments carried out in mouse and/or human cells (Figure 1b, Table S1).

**Figure 1.**
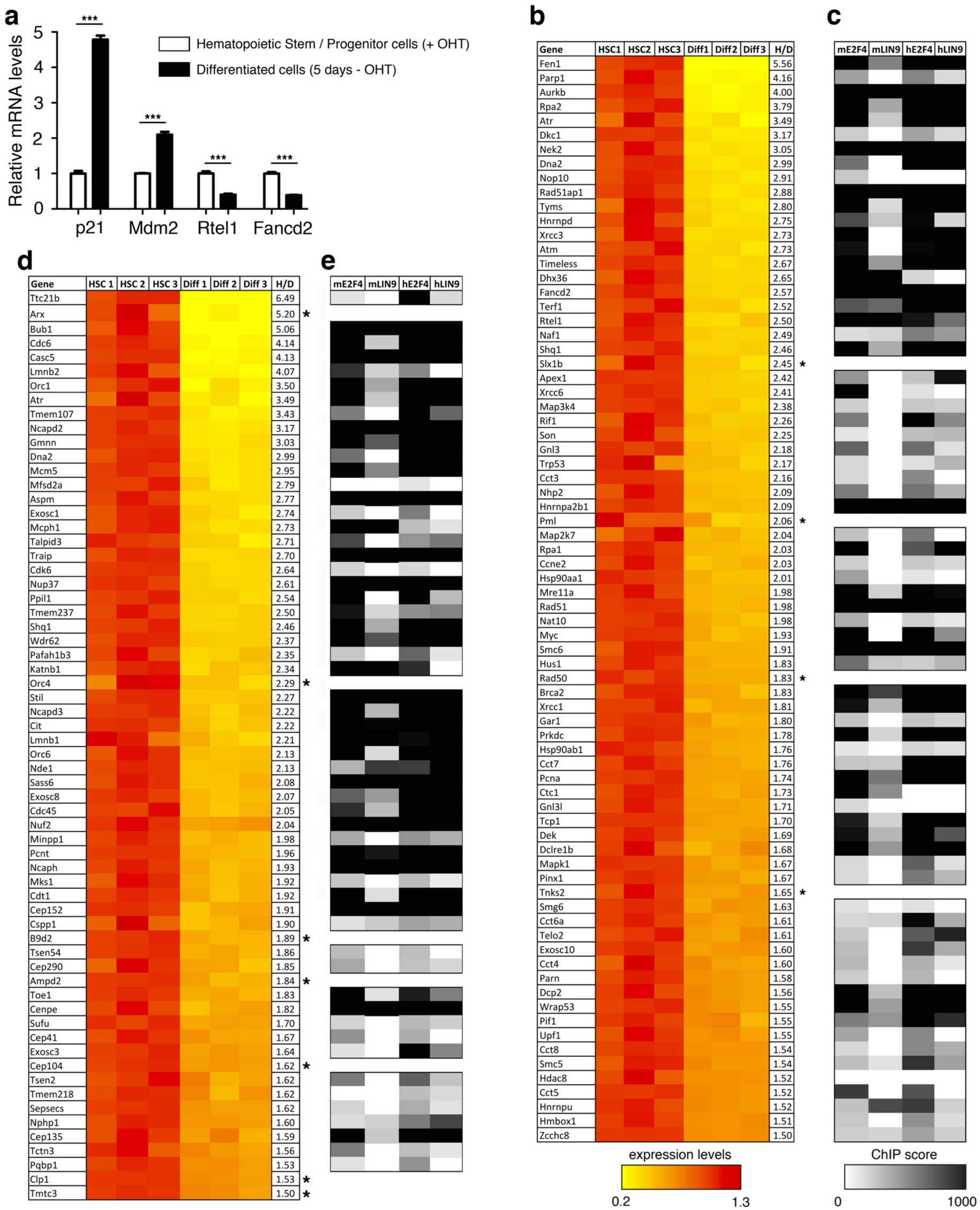
Telomere-related or microcephaly-related genes downregulated upon bone marrow cell differentiation, and their potential regulation by DREAM. **(a)** The differentiation of Hoxa9-ER expressing bone marrow cells (BMCs) correlates with p53 activation. Robust-multi average values for *p21/Cdkn1a*, *Mdm2*, *Rtel1* and *Fancd2* expression were extracted from transcriptome data of Hoxa9-ER expressing hematopoietic stem and progenitor cells grown in the presence of tamoxifen (OHT), or 5 days after tamoxifen withdrawal in differentiated cells; average values (from triplicates) in cells with tamoxifen were given a value of 1. Means + s.e.m. are shown; ***P ≤ 0.001 by Student’s t test. **(b)** Telomere-related genes downregulated upon the differentiation of Hoxa9-ER bone marrow cells. Expression values for 76 telomere-related genes, from triplicates; average values in cells with tamoxifen were given a value of 1. Genes are listed according to decreasing repression fold (H/D). According to the Target gene regulation database, 72/76 genes are downregulated upon mouse and/or human p53 activation (the 4 exceptions are mentioned by an asterisk). **(c)** For each of the 72 p53-regulated genes in (b), the highest chromatin immunoprecipitation (ChIP) scores of E2F4 or LIN9 binding in mouse (m) or human (h) cells are represented. Values are according to ChIP-Atlas. **(d)** Genes associated with microcephaly syndromes and downregulated upon BMC differentiation. Expression values and ChIP scores represented as in (b), for 64 genes associated with syndromes of microcephaly or cerebellar hypoplasia, of which 57 are downregulated by p53. **(e)** ChIP scores of E2F4 or LIN9 binding in mouse (m) or human (h) cells for the 57 microcephaly-related, p53-downregulated genes, represented as in (c).

**Table 1.** Genes associated with telomere-related ontology terms are over-represented among genes downregulated in differentiated bone marrow cells. Telomere-related gene ontology (GO) terms and descriptions are mentioned, as well as enrichment factors and P- and FDR q-values. As an example, genes with the GO term #0032204 (regulation of telomere maintenance) represent 80/17461 genes associated with a GO term, but 43/3631 of genes associated with a GO term and downregulated at least 1.5 times upon BMC differentiation, which represents a 2.58-fold enrichment.

| GO term | Description | P-value | FDR q-value | Enrichment (N, B, n, b) |
| --- | --- | --- | --- | --- |
| GO : 0032204 | regulation of telomere maintenance | 8.24E-11 | 1.1E-8 | 2.58 (17461,80,3631,43) |
| GO : 1904356 | regulation of telomere maintenance via telomere lengthening | 1.83E-9 | 1.98E-7 | 2.74 (17461,58,3631,33) |
| GO : 0032200 | telomere organization | 1.85E-9 | 1.99E-7 | 2.58 (17461,69,3631,37) |
| GO : 0000723 | telomere maintenance | 1.85E-9 | 1.98E-7 | 2.58 (17461,69,3631,37) |
| GO : 0032206 | positive regulation of telomere maintenance | 4.96E-8 | 4.36E-6 | 2.69 (17461,50,3631,28) |
| GO : 0032210 | regulation of telomere maintenance via telomerase | 1.35E-7 | 1.07E-5 | 2.65 (17461,49,3631,27) |
| GO : 1904358 | positive regulation of telomere maintenance via telomere lengthening | 4.89E-7 | 3.47E-5 | 2.89 (17461,35,3631,21) |
| GO : 0010833 | telomere maintenance via telomere lengthening | 1.1E-6 | 7.22E-5 | 3.54 (17461,19,3631,14) |
| GO : 0070203 | regulation of establishment of protein localization to telomere | 1.34E-6 | 8.43E-5 | 4.37 (17461,11,3631,10) |
| GO : 0007004 | telomere maintenance via telomerase | 1.59E-6 | 9.79E-5 | 4.07 (17461,13,3631,11) |
| GO : 1904851 | positive regulation of establishment of protein localization to telomere | 5.87E-6 | 3.25E-4 | 4.33 (17461,10,3631,9) |
| GO : 0032212 | positive regulation of telomere maintenance via telomerase | 3.16E-5 | 1.47E-3 | 2.64 (17461,31,3631,17) |
| GO : 0032205 | negative regulation of telomere maintenance | 2.29E-4 | 8.29E-3 | 2.34 (17461,35,3631,17) |

We previously showed that p53 activation leads to an increased binding of the E2F4 repressor at the promoter of 4 of these 72 telomere-related genes (*Brca2*, *Fancd2*, *Rad51*, *Rtel1*) [25], which contributed to provide evidence that p53-mediated gene repression often occurs indirectly, through the recruitment of the E2F4-containing complex DREAM, often close to transcription start sites. Thus, we used ChIP-Atlas [38] to search for evidence of E2F4 binding at the promoters of the 72 telomere-related, p53-regulated candidate genes we had identified. The data compiled from 18 ChIP-seq experiments revealed E2F4 binding at 71 out of the 72 genes, in regions frequently overlapping transcription start sites (Figure 1c, Table S2). To further identify candidate DREAM targets, we used ChIP-Atlas to search for evidence of MuvB binding. ChIP-Atlas does not provide information on LIN54, the DNA-binding subunit of MuvB, so we analyzed instead the ChIP-seq data with antibodies against LIN9, another subunit of MuvB. The data compiled from 4 ChIP-seq experiments indicated LIN9 binding at 36 out of the 72 genes, at regions overlapping the regions bound by E2F4 (Figure 1c, Table S2). Half the genes bound by E2F4 were not identified in LIN9 ChIP-seq experiments, which could suggest a regulation mediated by E2F4 independently of the DREAM complex. Alternatively, this might reflect technical limitations, resulting from the fact that LIN9 does not directly bind to DNA, or that the ChIP-seq data resulted from 18 experiments with antibodies against E2F4 but only 4 experiments with antibodies against LIN9, or from qualitative differences between the antibodies used in the experiments. The repertoires of genes downregulated by the p53-DREAM pathway appear to be well conserved between humans and mice [39], so we next analyzed ChIP-seq data from human cells. Evidence for binding by E2F4 was found for 68 out of the 72 homologous human genes, most often around the transcription start site, and 59 of these genes were also bound by LIN9 (Figure 1c, Table S3). Average ChIP-seq binding scores appeared slightly higher in human cells, particularly with antibodies against LIN9 (average score of 638 for 59 human genes, compared to 496 for 36 mouse genes). This might suggest that antibodies against LIN9 were more efficient to precipitate the human LIN9 protein, so that the number of murine genes downregulated by p53-DREAM was under-estimated due to technical difficulties. In sum, a total of 61 telomere-related genes were detected in ChIP assays with antibodies against E2F4 and LIN9 in at least one species (Figure 1c), strengthening the notion that the p53-DREAM pathway plays a significant role in regulating telomere biology.

We previously reported that p53 can also downregulate many genes of the Fanconi anemia DNA repair pathway, a pathway implicated in the repair of DNA interstrand cross-links [25]. Accordingly, GOrilla revealed a significant enrichment for genes of the “interstrand cross-link repair” GO term among the genes downregulated in murine differentiated BMCs (Table S4a). We found 55 genes down-regulated upon BMC differentiation, encompassing genes mutated in Fanconi anemia, regulating the Fanconi DNA repair pathway and/or belonging to the GOrilla interstrand cross-link repair GO term, or to a recently proposed list of FA-related genes [40], including 52 genes downregulated by p53 according to the TGR database (Figure S2a, Table S5). Out of these 52 genes, twelve are also known to impact on telomere biology (*Atm*, *Atr*, *Brca2*, *Dclre1b*, *Fancd2*, *Fen1, Hus1, Rad51, Rad51ap1, Rpa2, Telo2, Xrcc3*). ChIP-seq experiments revealed E2F4 binding at 51, and LIN9 binding at 41 of the 52 genes (Figure S2b), within regions frequently overlapping transcription start sites (Table S6). When we analyzed ChIP-seq data from human cells, evidence for binding by E2F4 was found for 51/52 homologous genes, and 52/52 genes were bound by LIN9 (Figure S2b) also at sequences frequently overlapping transcription start sites (Table S7).

Recent data suggested that some of the genes mutated in dyskeratosis congenita or Fanconi anemia may affect ribosomal function [41,42] and frameshift *TP53* mutations cause a pure red cell aplasia resembling Diamond-Blackfan anemia, together with relatively short telomeres [28]. We thus also determined if BMC differentiation altered the expression of genes involved in ribosome function. Indeed, among the genes downregulated at least 1.5-fold upon tamoxifen withdrawal, a significant enrichment was observed for 28 GO terms related to ribosome biology, rRNA biogenesis and maturation, or RNA polymerase I (Table S4b). These 28 GO terms are partially overlapping and correspond to 168 different genes, of which 10 (*Dkc1*, *Exosc10*, *Gar1*, *Gnl3l*, *Naf1*, *Nat10*, *Nhp2*, *Nop10*, *Prkdc*, *Shq1*) are also known to impact on telomere biology. Furthermore, we noticed 3 additional genes encoding subunits of RNA Polymerase I (*Polr1d*, *Taf1c*, *Taf1d*) that were downregulated upon BMC differentiation, raising the total of candidates to 171 genes, of which 162 are downregulated by p53 according to the TGR database (Figure S3, Table S8). E2F4 binding was found at 152, and LIN9 binding at 50 of the 162 murine genes, at sequences frequently overlapping transcription start sites (Figure S3, Table S9). ChIP-seq data from human cells indicated binding by E2F4 for 153/162, and by LIN9 for 115/162 homologous genes, often within regions overlapping transcription start sites (Figure S3, Table S10).

We next enquired if genes mutated in other bone marrow disorders might be downregulated at least 1.5 fold upon BMC differentiation, and found 17 candidate genes: *Ankrd26*, *Etv6* and *Mastl*, mutated in thrombocytopenia; *Pik3r1*, *Tcf3* and *Cd79b*, mutated in agammaglobulinemia; *Cdan1* and *Sec23b*, mutated in congenital dyserythropoietic anemia; *G6pc3* and *Gfi1*, mutated in severe congenital neutropenia; *Rbm8a*, mutated in the thrombocytopenia absent radius syndrome; *Efl1*, mutated in the Shwachman-Diamond syndrome type 2; *Rab27a*, mutated in the Griscelli syndrome type 2; *Mtr,* mutated in homocystinuria megaloblastic anemia; *Mthfd1,* mutated in combined immunodeficiency and megaloblastic anemia with or without hyperhomocysteinemia; *Dnajc21*, mutated in bone marrow failure syndrome 3; and *Nuf2*, mutated in a bone marrow failure syndrome with microcephaly and renal hypoplasia. Out of these 17 genes, 15 were reported to be downregulated upon p53 activation (Figure S4, Table S11). Out of the 15 genes, 14 were bound by E2F4 and 6 were bound by LIN9 in murine cells (Figure S4, Table S12) and 13 were bound by E2F4 and 12 by LIN9 in human cells (Figure S4, Table S13).

We also used the Human Phenotype Ontology website from the Jackson Laboratory (https://hpo.jax.org) [43] to search for genes associated with abnormalities of blood and blood-forming tissues (ontology term #HP:0001871), and found that, out of a list of 1322 genes, 336 candidates were downregulated at least 1.5 times upon murine BMC differentiation, including 277 reported to be downregulated by p53 according to the Target gene database (Figure S5, Table S14). Out of these 277 genes, 243 were bound by E2F4 and 102 by LIN9, close to the transcription start site in most cases (Figure S5, Table S15). Out of the 277 human homologous genes, 245 were bound by E2F4 and 198 by LIN9 (Figure S5, Table S16).

Together, the differentiation of bone marrow cells correlated with the decreased expression of a total of 571 genes implicated in hematopoiesis, including 499 genes downregulated by p53 according to the TGR database (Table S17a-b, see also Figure 3c for a summary of our approach). For 374 of these genes, E2F4 and LIN9 were found to bind at identical regions in at least one species (Table S17c, Figure 3c). Furthermore, to focus on the best candidate p53-DREAM targets, we also considered the ChIP scores for E2F4 and LIN9 binding for each of the 374 genes. For each gene, we added the ChIP scores of E2F4 and LIN9 in both species, for a maximal value of 4000 (Table S18). Total ChIP scores ranged from 313 to 4000, and we noticed total ChIP scores of 656 and 720, respectively for *Fbl* and *Dkc1,* two genes reported to be directly repressed by p53 binding [26,44] and a score of 979 for *Exosc5*, a gene previously proposed to be regulated by DREAM [14]. We thus considered the 269 genes with a total ChIP score ≥ 979 as the most likely candidate p53-DREAM targets (Tables S17d, S18 and Figure 3c).

**Figure 3.**
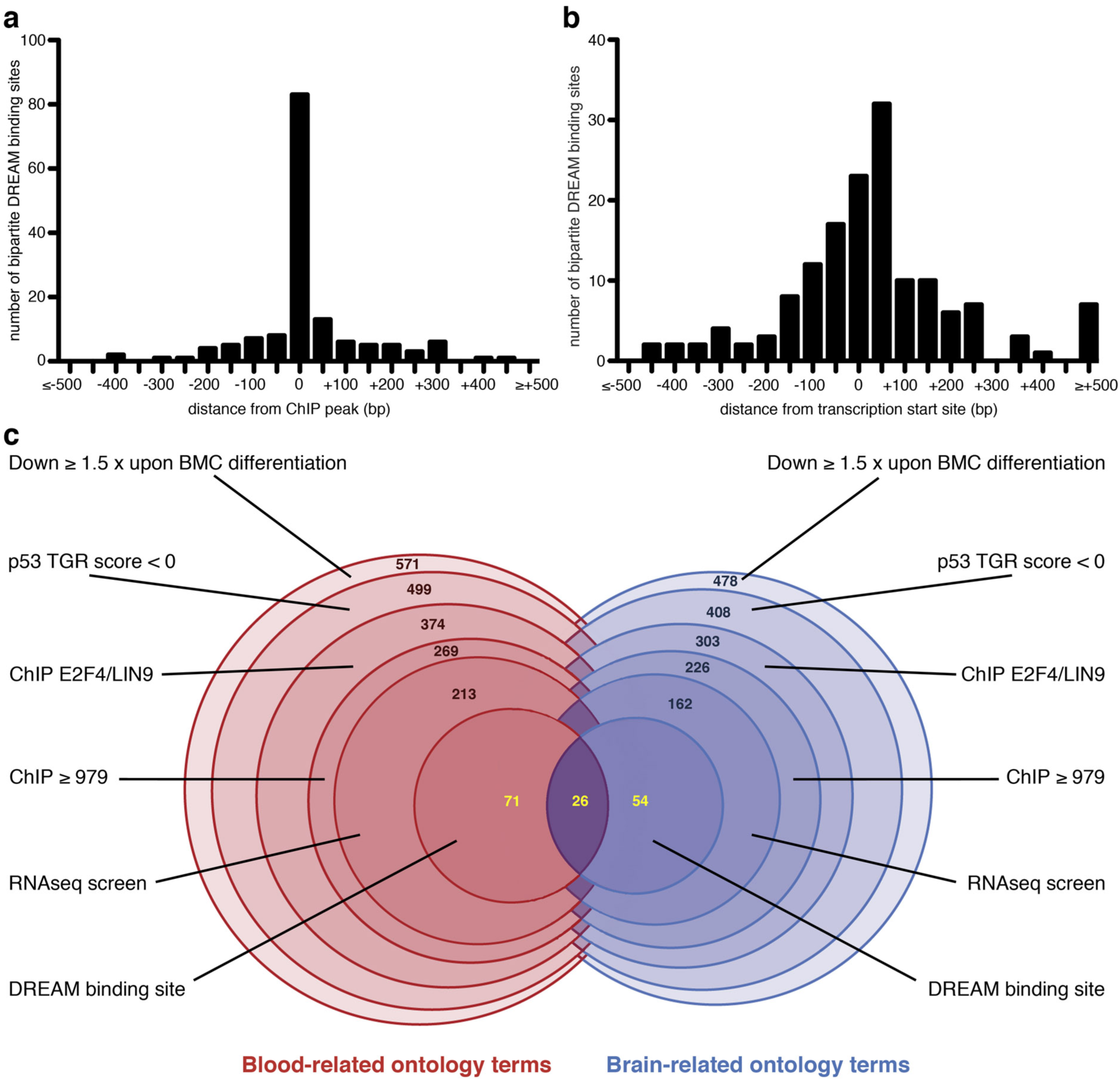
Mapping of putative DREAM binding sites and summary of our systematic approach. **(a)** Mapping of the putative DREAM binding sites relative to the ChIP peaks of E2F4 and/or LIN9 binding in 50 bp windows, for the 151 genes listed in Table 2. **(b)** Mapping of the putative DREAM binding sites relative to the transcription start sites of the 151 genes, in 50 bp windows. **(c)** Venn-like diagram of our systematic approach. Numbers in black indicate, at each step of the approach, genes related either to blood-related ontology terms or to brain-related ontology terms, which were analyzed separately (see text for details). In the last step, blood- and brain-related candidate genes were analyzed together to search for DREAM binding sites, and numbers in yellow indicate genes related to blood-related terms only, brain-related terms only, or both blood- and brain-related terms. Detailed listings of genes can be found in Tables S17 and S26.

**Table 2.**
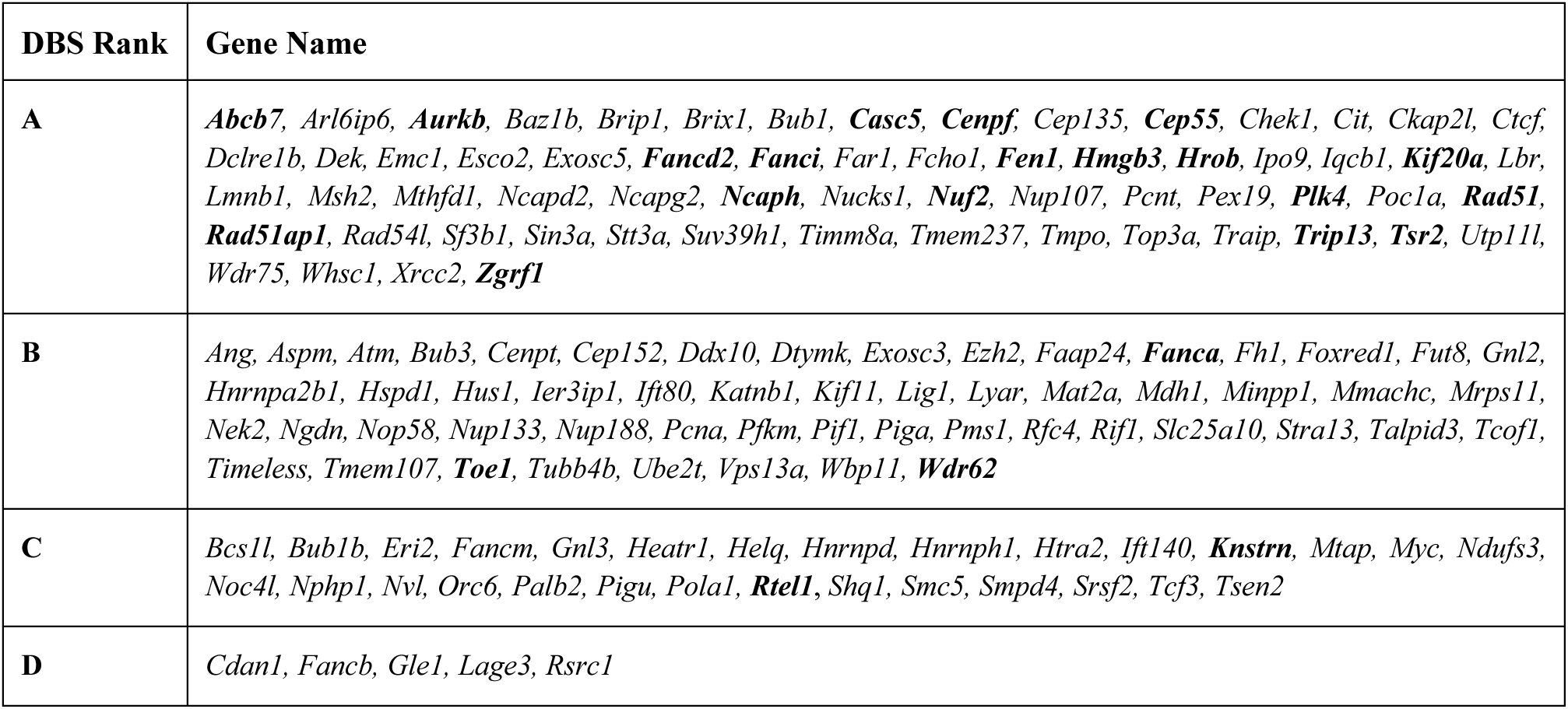
A summary of genes with appropriately mapped putative DREAM binding sites. For each gene, putative bipartite DREAM binding sites (DBS) were searched for in the regions bound by E2F4 and LIN9, by using a positional frequency matrix (PFM10 or PFM22). Putative DBS were identified for 151 candidate p53-DREAM target genes. A combination of PFM score and degree of DNA sequence conservation was used to classify candidate DBS into 4 categories, with A corresponding to the better candidates (see Tables S29-S34 for details). If a same DBS was found with PFM10 and PFM22, the ranking with PFM22 is mentioned here. For genes with several putative DBS, only the experimentally tested (in bold) or the highest-ranking element is mentioned here (all putative DBS are included in supplementary tables). In the present study, 21 DBS were tested in luciferase assays.

To estimate the relevance of this list of 269 candidates, we analyzed the dataset GSE171697, which includes RNAseq data from hematopoietic stem cells of unirradiated p53 KO mice, unirradiated WT mice, or irradiated WT mice [45]. We also analyzed GSE204924, with RNAseq data from splenic cells of irradiated p53^Δ24/-^ or p53^+/-^ mice [46]. The public data from this dataset, although incomplete, appeared interesting because p53^Δ24^ is a mouse model prone to bone marrow failure [47] and the spleen is an hematopoietic organ in mice [48]. As expected, increased p53 activity correlated with an average increase in expression for 15 genes known to be transactivated by p53, with 13/15 genes upregulated at least 1.5-fold (Table S19). By contrast, only 56/269 candidate p53-DREAM targets genes appeared upregulated in cells with increased p53 activity (Table S19). These 56 genes were considered poor candidate p53-DREAM targets and removed from further analyses, leading to a list of 213 candidate p53-DREAM targets related to blood abnormalities (Tables S17e, S19 and Figure 3c).

### Candidate p53-DREAM target genes associated with brain abnormalities

We next searched for p53-DREAM target genes whose altered expression might contribute to brain abnormalities. Many genes whose mutations cause microcephaly or cerebellar hypoplasia encode proteins implicated in fundamental processes common to all somatic cells (e.g. chromosome condensation, mitotic spindle activity, tRNA splicing). We thus reasoned that BMC differentiation data might also be exploited to search for genes downregulated upon p53 activation and implicated in these diseases. To test this, we used a candidate approach and searched for genes that might be regulated by the p53-DREAM pathway among the 30 genes mutated in primary microcephaly, 23 genes mutated in pontocerebellar hypoplasia, 39 genes mutated in hypoplasia of the cerebellar vermis (Joubert syndrome), 18 genes mutated in syndromes combining microcephaly and dwarfism (Seckel syndrome, Meier-Gorlin syndrome, microcephalic osteodysplastic primordial dwarfism), 12 genes mutated in lissencephaly (often associated with microcephaly); *Nuf2*, mutated in a bone marrow failure syndrome with microcephaly and renal hypoplasia; *Pafah1b3*, truncated in a case of brain atrophy*; Pqbp1*, mutated in the Renpenning syndrome (a X-linked syndrome of microcephaly); and *Shq1*, mutated in a syndrome with cerebellar hypoplasia, dystonia and seizures; for a total of 126 candidate genes. A downregulation of gene expression of at least 1.5-fold upon BMC differentiation was found for 64 of these candidates, including 57 reported to be downregulated upon p53 activation according to the TGR database (Figure 1d, Table S20). Out of the 57 genes, 55 were bound by E2F4 and 36 by LIN9, within regions overlapping the transcription start site in most cases (Figure 1e, Table S21). Out of the 57 human homologs, all were bound by E2F4 and 49 by LIN9 (Figure 1e, Table S22).

We next searched the Human Phenotype Ontology website from Jackson Laboratory for genes associated with microcephaly or cerebellar hypoplasia (ontology terms #HP:0000252 and HP:0007360) and found that, out of a list of 1430 genes, 474 candidates were downregulated at least 1.5 times upon murine BMC differentiation, including 404 reported to be downregulated upon p53 activation (Figure S6, Table S23). Out of these 404 genes, 354 were bound by E2F4 and 153 by LIN9, in regions overlapping transcription start sites in most cases (Figure S6, Table S24). Out of the 404 human homologous genes, 371 were bound by E2F4 and 292 by LIN9 (Figure S6, Table S25).

In sum, the differentiation of bone marrow cells correlated with the decreased expression of 478 genes implicated in microcephaly or cerebellar hypoplasia, including 408 downregulated upon p53 activation according to the TGR database (Table S26a-b and Figure 3c). For 303 of these genes, E2F4 and LIN9 were found to bind at identical regions in at least one species (Table S26c, Figure 3c). Furthermore, total ChIP scores ≥ 979 were found for 226 of the 303 genes, which appeared as better candidate p53-DREAM targets (Tables S26d, S27, Figure 3c).

To estimate the relevance of this list of 226 candidates, we analyzed datasets GSE78711 and GSE80434, containing RNAseq data from human cortical neural progenitors infected by the Zika virus (ZIKV) or mock-infected, because ZIKV was shown to cause p53 activation in cortical neural progenitors and microcephaly [49,50]. Accordingly, most genes (12/16) known to be transactivated by p53 were upregulated in ZIKV-infected cells (Table S28). By contrast, only 64/226 candidate p53-DREAM targets genes appeared upregulated in ZIKV-infected cells (Table S28). These 64 genes were considered poor candidate p53-DREAM targets and removed from further analyses, leading to a list of 162 candidate p53-DREAM targets related to brain abnormalities (Tables S26e, S28 and Figure 3c). Importantly, out of the 162 microcephaly-related candidate genes identified (Table 26e), 58 also belonged to the list of 213 genes associated with abnormal hematopoiesis (Table S17e), consistent with the notion that a deregulation of the p53-DREAM pathway might be involved in both pathological processes. In sum, we identified 317 genes (213 + 162 - 58) downregulated upon bone marrow cell differentiation and p53 activation, bound by E2F4 and LIN9 in at least one species, with total ChIP scores ≥ 979, and which appeared as better candidate p53-DREAM targets after analyzing appropriate RNAseq data (Tables S17, S26).

### Identification of DREAM binding sites in candidate target gene promoters

We aimed to obtain further evidence of a DREAM-mediated regulation for the better candidates by searching for putative DREAM binding sites (DBS) within the regions bound by E2F4 and/or LIN9. Among the 213 candidate genes associated with blood abnormalities found here, we previously identified well-conserved bipartite DBS, functional in both mouse and human species, for *Fancd2, Fanci* and *Rad51* [25]. Accordingly, we next used DNA sequence conservation as a criterion to identify the best putative bipartite DBS within the regions bound by E2F4 and LIN9. We created a positional frequency matrix based on 10 functionally demonstrated murine DBS (PFM10, Figures 2a and S7a) and used PWMScan [51] to search for putative DBS in both mouse (mm10) and human (hg38) genomes, with a P-value threshold of 10^-3^. Based on our previous data with *Fanc* genes [25], we focused our search on DBS in the same orientation as the gene transcripts. This led to identify putative DBS for 55 genes associated with blood abnormalities (Table S29). A combination of PFM score and degree of DNA sequence conservation was used to classify candidate DBS into 4 categories: ranks A-C for DBS with positive PFM scores and respectively 0-1, 2-3 or 4 mismatches between mouse and human sequences at positions 2-6 or 11-16 of the consensus sequence, and rank D for DBS with negative PFM scores and 0-1 mismatch (see Table S29 for details). Likewise, we used PWMScan with PFM10 and sequence conservation to identify putative DBS in promoters of the 162 genes associated with brain abnormalities. DBS with various PFM score and DNA sequence conservation were identified for 52 genes, of which 15 were also associated with blood abnormalities (Table S30).

**Figure 2.**
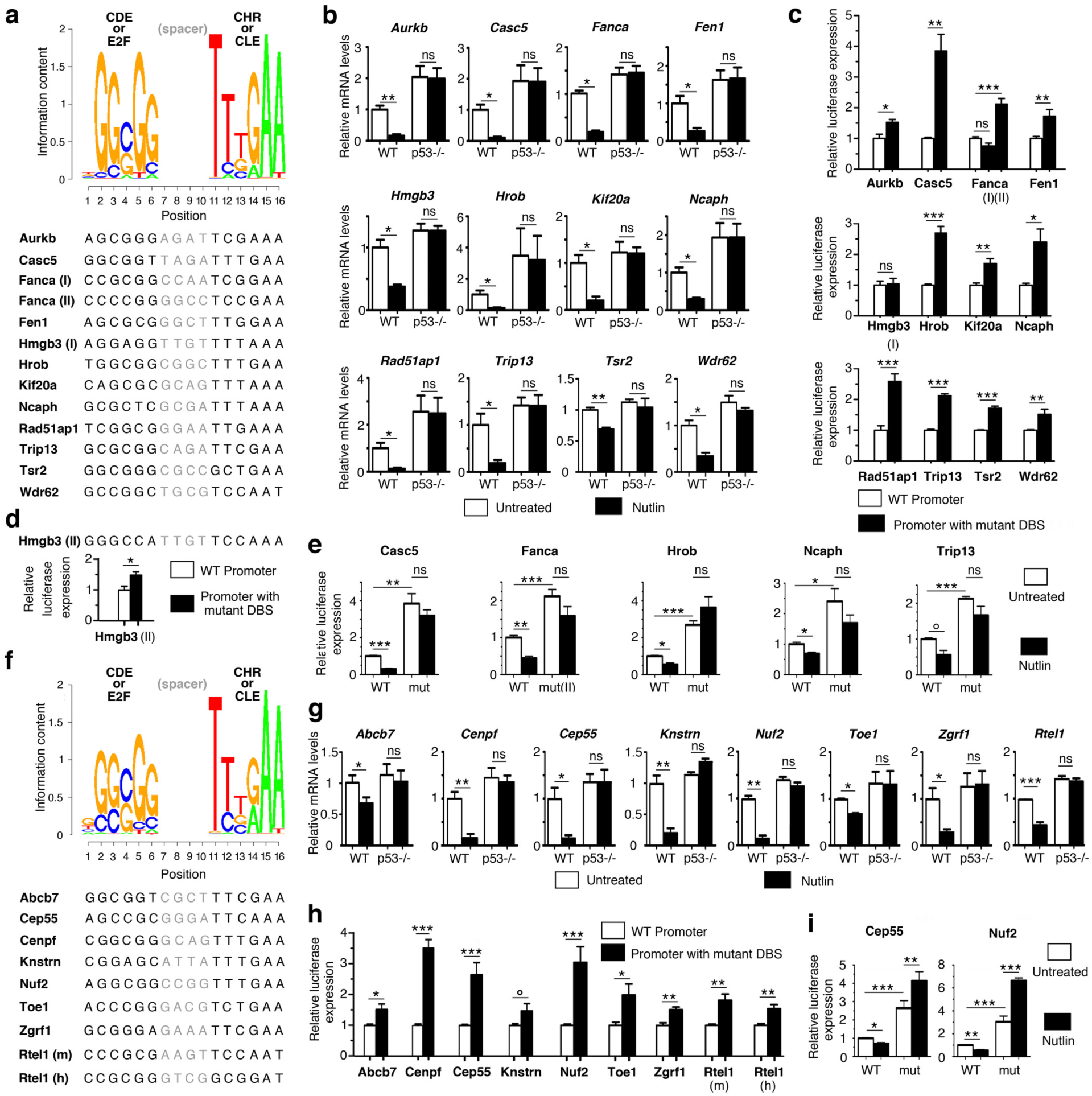
Functional assays of putative DREAM binding sites identified with positional frequency matrices. **(a)** Logo of the positional frequency matrix PFM10 and DNA sequence of 13 murine putative DREAM binding sites (DBS) identified with this matrix. PFM10 results from the DNA sequence of 10 experimentally validated DBS. Spacer DNA sequences between the GC-rich (CDE or E2F) and AT-rich (CHR or CLE) elements were not used to define the matrix (see Figure S7a for details). **(b)** In mouse embryonic fibroblasts (MEFs), p53 activation leads to the downregulation of the 12 tested genes. mRNAs from wild-type (WT) and p53^-/-^ MEFs, untreated or treated with 10 mM Nutlin (a MDM2 antagonist) for 24h, were quantified using real-time PCR, normalized to control mRNAs, then the amount in WT untreated cells was assigned a value of 1. Means + s.e.m. from 3 independent experiments are shown. Similar results for *Fanca* and *Fen1* were reported previously [25]. **(c)** Luciferase assays of the putative DBS reported in (a). For each candidate gene, a 1-1.5 kb fragment containing WT promoter sequences or the same promoter with point mutations affecting the DBS was cloned upstream a luciferase reporter gene (tested DBS were from the species with best PFM score). WT or mutant luciferase reporter plasmids were transfected into NIH3T3 cells, luciferase activity was measured 24h after transfection, and activity with the WT construct was assigned a value of 1. Results from ≥ 4 values and ≥ 2 independent cellular experiments for each tested plasmid. **(d)** DNA sequence of an alternate putative DBS at the *Hmgb3* promoter and its validation by luciferase assays, carried out as above. **(e)** Mutating the DBS at the *Casc5*, *Fanca*, *Hrob*, *Ncaph* or *Trip13* promoter abrogates its p53-dependent repression. WT or mutant (mut) luciferase plasmids were transfected into NIH3T3 cells, untreated or treated with Nutlin, then luciferase activity was measured after 24h. Results from ≥ 4 values and ≥ 2 independent cellular experiments for each tested plasmid. **(f)** Logo of PFM22 and DNA sequence of 9 tested DBS. PFM22 results from 10 DBS validated in previous reports and 12 DBS tested in the present study, with spacer DNA sequences not used to define the matrix (see Figure S7b for details). **(g)** In MEFs, p53 activation leads to the downregulation of the 8 tested genes. mRNAs from WT and p53^-/-^ MEFs were treated and analyzed as in (b). Similar results for *Rtel1* were reported previously [26]. **(h)** Luciferase assays of the putative DBS reported in (f), performed as in (c). **(i)** Mutating the DBS at the *Cep55* or *Nuf2* promoter abrogates its p53-dependent repression. Luciferase assays were performed as in (e). \*\*\**P<*0.001, \*\**P<*0.01, \**P<*0.05, °*P*≤0.07 in Student’s *t* tests.

A fraction of the putative DBS identified with this approach were already shown to be functional in previous reports by using luciferase assays. This is the case for at least one of the two overlapping DBS at the human *Aurkb* promoter [52], the murine DBS at the *Plk4* promoter [53], and for both murine and human DBS at the *Fancd2*, *Fanci* and *Rad51* promoters [25] (Tables S29, S30). Although testing the functionality of all the putative DBS was beyond the scope of our study, we aimed to test the validity of our predictions by performing luciferase assays on a subset of the elements. We tested the putative DBS of the 12 following genes: *Hmgb3*, *Hrob*, *Ncaph* and *Trip13*, containing a putative DBS of rank A; *Aurkb*, containing two overlapping DBS of ranks A and B (shifted by only 1 nucleotide and thus similar to a single DBS); *Fanca*, containing two non-overlapping putative DBS of ranks B and C; *Wdr62*, containing a putative DBS of rank B; and *Casc5*, *Fen1*, *Kif20a*, *Rad51ap1* and *Tsr2*, containing putative DBS of rank D (Figure 2a, Tables S29, S30). According to gene ontology, these genes are associated with either abnormal hematopoiesis (*Aurkb*, *Fen1*, *Hrob*, *Kif20a*, *Rad51ap1*, *Tsr2*), microcephaly (*Casc5*, *Hmgb3*, *Ncaph, Wdr62*), or both (*Fanca*, *Trip13*). For genes associated with abnormal hematopoiesis, we first verified that their expression was decreased in BMCs from p53^Δ31/Δ31^ mice, prone to bone marrow failure, compared to WT BMCs (Figure S8). We next determined, as a prerequisite to luciferase assays, that the expression of all tested genes, as well as their p53-mediated repression, could be observed in mouse embryonic fibroblasts (MEFs), because luciferase assays rely on transfections into the mouse embryonic fibroblast cell line NIH3T3 (Figure 2b). We cloned the promoters of the candidate targets upstream of a luciferase reporter gene, then introduced point mutations specific to the putative DBS element to abolish its potential function. In these experiments, the DBS for murine *Aurkb* served as a positive control because of its high sequence conservation with the DBS shown to be functional in the homologous human gene [52]. Consistent with its expected role in gene repression, the mutation of the DBS for murine *Aurkb* led to increased luciferase expression (Figure 2c). A similar result was obtained with DBS at 10/11 other tested genes (*Casc5*, *Fanca* (DBS II), *Fen1*, *Hrob*, *Kif20a*, *Ncaph*, *Rad51ap1*, *Trip13*, *Tsr2, Wdr62*; Figure 2c). For *Hmgb3* however, the putative DBS element did not appear to be functional in luciferase assays. We reasoned that an improved positional frequency matrix, that would include the 11 additional DBS we had just tested might lead to identify a proper DBS for this gene. Indeed, the second matrix (PFM21) suggested a new putative DBS at the *Hmgb3* promoter, whose mutation affected gene expression in luciferase assays (Figure 2d, Table S30). Of note, NIH3T3 cells exhibit an attenuated p53 pathway compared to primary MEFs (Figure S9a). This facilitates cell survival after lipofections required in luciferase assays but leads to decreased p53-DREAM-mediated gene repression (Figure S9b). Under these experimental conditions, p53 activation in transfected NIH3T3 cells led to the robust repression (>1.4 fold) of 5 WT promoters cloned upstream the luciferase reporter gene (the promoters for *Casc5*, *Fanca*, *Hrob*, *Ncaph* and *Trip13*). Importantly, the p53-mediated repression of these 5 promoters was abrogated by mutating the identified DBS (Figure 2e), providing direct evidence of the functional relevance of DBS identified with our positional frequency matrix.

These experiments indicated that we could identify sites impacting on luciferase expression for 12/12 tested genes, and we next integrated these sites into a third positional frequency matrix (PFM22), used in all further analyses (Figure 2f and S7b). We used PWMscan with PFM22 and a P-value threshold of 10^-3^ to reanalyze the genes for which putative DBS had been suggested by using PFM10 (Tables S31, S32). We reasoned that good candidate DBS identified with PFM10 were likely to be found again with PFM22: this was verified for 45/55 hematopoiesis-related genes and 37/52 microcephaly-related genes. Furthermore, alternative DBS (often with better scores) were suggested with PFM22 for 7/55 hematopoiesis-related genes and 9/52 microcephaly-related genes. For a few genes (e.g. *Cdan1*, *Gle1*), the putative DBS identified with PFM10 were not detected with PFM22 and appeared as potentially weaker candidates. We also considered the converse situation - that for some genes for which no DBS had been suggested with PFM10, it might be possible to find putative DBS with PFM22. Indeed, the use of PFM22 made it possible to find a putative DBS for 57 additional targets (Table S33).

Thus, out of 317 genes associated with blood and/or brain abnormalities that appeared as potential DREAM targets, we found 149 genes containing at least one appropriately mapped putative bipartite DBS, in the same orientation as transcription, and with partial or complete DNA sequence conservation between human and mouse. These genes include *Abcb7*, *Cep55*, *Cenpf*, *Knstrn*, *Nuf2*, *Toe1* and *Zgrf1*, for which putative DBS were also tested in luciferase assays (Figure 2g-i). As for the genes for which no DBS was suggested with our PFMs, we hypothesized that they might be regulated via DBS not fulfilling our criteria. For example, a CDE/CHR was shown to regulate the expression of murine *Ccnb2*, but only the CHR element was conserved in the homologous human gene [54,55]. Similarly, we identified a bipartite DBS in the murine *Rtel1* promoter for which only the CDE (E2F) element was conserved in the human homolog, and a DBS in the human *RTEL1* promoter for which only the CDE (E2F) element was conserved in the murine homolog (Figure 2g-h and Table S34). Similar cases, *i.e.* putative DBS with a positive score in one species and a perfect conservation of either the CDE (E2F) at positions 2-6, or the CHR (CLE) at positions 11-16, were found for 5 other genes (Table S34). Of note, because these sites correspond to DBS with positive scores and limited DNA sequence conservation, most had already been detected as sites of rank C (at the promoters of *Helq*, *Htra2*, *Ndufs3* and *Smc5*). Accordingly, DBS with positive PFM22 scores and either 4 mismatches anywhere in the DBS or more mismatches but affecting only the CDE (E2F) or only the CHR (CLE) were together classified as rank C sites. Finally, our PFMs were designed to identify bipartite DBS with a CDE (E2F) motif separated from a CHR (CLE) motif by a spacer of 4 bp. Presumably, candidate DREAM targets for which no DBS was identified with these PFMs might be bound by DREAM either via a bipartite site with spacer sequences of a different length, or by a single E2F or a single CHR motif, as previously proposed [14,22].

Table 2 summarizes our results: putative DREAM-binding sites were identified at the promoter of 151 genes, including 97 genes associated with blood ontology terms and 80 with brain ontology terms (Tables S17f and S26f, Figure 3c). Consistent with a functional relevance of the predicted DBS, most sites co-mapped with peaks of E2F4 and/or LIN9 binding (Figure 3a). At the 151 promoters, 83 putative DBS mapped in a 50 bp-long window centered on ChIP peaks (Figure 3a), whereas the frequency of putative DBS per 50 bp-long window was 4 10^-4^ over the entire human genome, indicating a 1300-fold enrichment of DBS at ChIP peaks. This significant enrichment (f=3 10^-239^ in a hypergeometric test) is most likely underestimated because mouse-human DNA sequence conservations were not determined for putative DBS over the full genome. In addition, it was proposed that DREAM primarily associates with nucleosomes near the transcription start sites of its targets [56] and the distribution of predicted DBS was consistent with this notion (Figure 3b). Altogether, the differentiation of bone marrow cells correlated with the downregulation of 571 genes associated with blood-related ontology terms and 478 genes associated with brain-related ontology terms (Figure 3c), for a total of 883 genes (166 genes being associated with both blood- and brain-related terms, see Tables S17a and S26a). Out of these 883 genes, 760 (499 + 408 – 147) were reported to be downregulated by p53 (Figure 3c, Tables S17b and S26b). Among those genes, our systematic approach identified 317 likely p53-DREAM targets, and our positional frequency matrices appeared as powerful tools to predict DREAM binding sites for about half of these target genes (Figure 3c, Tables S17e-f and S26e-f).

## Discussion

The capacity of p53 to activate the transcription of many targets, including genes important for cell cycle arrest (e.g. *CDKN1A*), apoptosis (e.g. *BAX*, *PUMA*) or cellular metabolism (e.g. *TIGAR*), has been recognized for decades. On the opposite, the potential importance of p53-dependent transcriptional repression has only emerged in recent years, in part because the mechanisms underlying p53-mediated repression remained controversial. In this report, we provide evidence for a general role of the p53-DREAM pathway in regulating genes associated with blood and/or brain abnormalities. We identified 317 potential p53-DREAM targets, *i.e.* genes with a decreased expression associated with murine bone marrow cell differentiation and p53 activation, and whose promoter sequences can be significantly bound by two subunits of the DREAM complex in mouse and/or human cells. Among these potential targets we identified putative DREAM binding sites in the promoter of 151 genes, and the mutation of a subset of these binding sites affected gene expression in luciferase assays.

Our approach has methodological similarities with the approaches described by Fischer *et al.*, who first provided evidence that p53 often represses transcription indirectly via the DREAM or RB/E2F pathways [10], then reported lists of most likely candidate p53-DREAM targets - a first list of 210 genes, most of which were regulators of the G2/M phases of the cell cycle [12], then a list of 971 G1/S or G2/M cell cycle genes [14]. Here, we found 883 genes related to blood and/or brain ontology terms downregulated upon BMC differentiation, of which 760 were reported to be downregulated by p53. E2F4 and LIN9 were found to bind at the promoters of at least 317 genes downregulated by p53, consistent with a major role of the DREAM complex in p53-mediated repression. Interestingly however, out of the 151 p53-DREAM targets with putative DBS we identified, only 30 were in the first list of 210 candidate DREAM targets, and 95 in the second list of 971 candidate DREAM targets reported by Fischer *et al.* The differences in p53-DREAM target repertoires might result in part from the fact that Fischer *et al.* mostly analyzed human fibroblasts treated with doxorubicin or nutlin, whereas we analyzed the effects of murine bone marrow cell differentiation. Interestingly, we identified *Brip1* as a p53-DREAM target gene, downregulated upon the differentiation of bone marrow cells (Figure S2) and in Zika-infected neural progenitors (Table S28), but not upon irradiation of hematopoietic stem cells (Table S19), consistent with the notion that different cellular responses might regulate partially distinct repertoires of DREAM targets. In addition, compared to Fischer *et al.*, our systematic use of pathology-related gene ontology likely created a sharper focus on clinically relevant target genes. In support of this, the list of 210 genes by Fischer *et al.* included only one gene mutated in Fanconi anemia (*Fancb*) and no gene mutated in dyskeratosis congenita [12], whereas in a previous study with mouse fibroblasts focusing on these bone-marrow failure syndromes [25] we found evidence for the p53-mediated repression of 8 clinically relevant genes that belong to our current list of 151 targets (*Fanca*, *Fancb*, *Fancd2*, *Fanci*, *Palb2*, *Rad51*, *Rtel1* and *Ube2t*).

Cells with a knock-out of Lin37, a subunit of the DREAM complex, can also be used to identify potential DREAM targets [16,17]. For example, Mages *et al.* used CRISPR-Cas9 to generate Lin37 KO murine cells, which were then rescued by an episomal Lin37 expression vector, and Lin37 KO and Lin37-rescued cells were compared by RNAseq analyses [17]. Our list of 151 genes overlaps only partially with the list of candidate DREAM targets obtained with this approach, with 51/151 genes reported to be downregulated in Lin37-rescued cells [17]. To better evaluate the reasons for this partial overlap, we extracted the RNAseq data from Lin37 KO and Lin37-rescued cells and focused on the 151 genes in our list. For the 51 genes that Mages *et al.* reported as downregulated in Lin37-rescued cells, an average downregulation of 14.8-fold was observed (Figure S10, Table S35). Furthermore, when each gene was tested individually, a downregulation was observed in all cases, statistically significant for 47 genes, and with a P value between 0.05 and 0.08 for the remnant 4 genes (Table S35). By contrast, for the 100 genes not previously reported to be downregulated in Lin37-rescued cells, an average downregulation of 4.7-fold was observed (Figure S10, Table S35), and each gene appeared downregulated, but this downregulation was statistically significant for only 35/100 genes, and P values between 0.05 and 0.08 were found for 23/100 other genes (Table S35). These comparisons suggest that, for the additional 100 genes, a more subtle decrease in expression, together with experimental variations, might have prevented the report of their DREAM-mediated regulation in Lin37-rescued cells.

Importantly, our approach integrated evolutive positional frequency matrices to identify putative bipartite DBS in the promoters of candidate target genes. Most putative DBS co-mapped with ChIP peaks for DREAM subunits and transcription start sites (TSS), and most DBS tested experimentally were found to affect gene expression in luciferase assays, suggesting reliable DBS predictions. The Target gene regulation (TGR) database of p53 and cell-cycle genes was reported to include putative DREAM binding sites for human genes, based on separate genome-wide searches for 7 bp-long E2F or 5 bp-long CHR motifs [23]. We analyzed the predictions of the TGR database for the 151 genes for which we had found putative bipartite DBS. A total of 342 E2F binding sites were reported at the promoters of these genes, but only 64 CHR motifs. The similarities between the predicted E2F or CHR sites from the TGR database and our predicted bipartite DBS appeared rather limited: only 14/342 E2F sites overlapped at least partially with the GC-rich motif of our bipartite DBS, while 27/64 CHR motifs from the TGR database exhibited a partial overlap with the AT-rich motif. Importantly, most E2F and CHR sites from the TGR database mapped close to E2F4 and LIN9 ChIP peaks, but only 16% of E2Fs (54/342), and 33% of CHRs (21/64) mapped precisely at the level of these peaks (Figure S11), compared to 55% (83/151) of our bipartite DBS (Figure 3a). Thus, at least for genes with bipartite DREAM binding sites, our method relying on PFM22 appeared to provide more reliable predictions of DREAM binding than the E2F and CHR sites reported separately in the TGR database. Importantly however, predictions of the TGR database may include genes regulated by a single E2F or a single CHR that would most likely remain undetected with PFM22, suggesting that both approaches provide complementary results. Of note, we previously used Consite (http://consite.genereg.net) [57] with positional frequency matrices (PFMs) from 6 or 8 experimentally demonstrated murine DBS [25,58] to search for bipartite DBS, a method suitable for the analysis of small (≤10 kb) DNA sequences. Here, the use of PWMscan with PFMs from 10 or 22 DBS made it possible to perform genome-wide searches for bipartite DBS, while facilitating the comparison of mouse and human DNA sequences. Our improved approach notably led to identify a functional DBS for *Fanca*, a gene we previously found downregulated by p53 but for which a DBS remained to be identified [25,26].

Finding a functionally relevant DBS for *Fanca,* mutated in 60% of patients with Fanconi anemia [59,60], may help to understand how a germline increase in p53 activity can cause defects in DNA repair. Importantly however, we previously showed that p53^Δ31/Δ31^ cells exhibited defects in DNA interstrand cross-link repair, a typical property of Fanconi anemia cells, that correlated with a subtle but significant decrease in expression for several genes of the Fanconi anemia DNA repair pathway, rather than the complete repression of a single gene in this pathway [25]. Thus, the Fanconi-like phenotype of p53^Δ31/Δ31^ cells most likely results from a decreased expression of not only *Fanca*, but also of additional p53-DREAM targets mutated in Fanconi anemia such as *Fancb*, *Fancd2*, *Fanci*, *Brip1*, *Rad51*, *Palb2*, *Ube2t* or *Xrcc2*, for which functional or putative DBS were also found with our systematic approach. Furthermore, our identification of DBS for *Rtel1*, a gene mutated in 30% of patients with the Hoyeraal-Hreidarsson syndrome [61–64] and whose expression correlated with the survival of p53^Δ31/Δ31^ mice [26], might help to explain how a germline increase in p53 activity can cause defects in telomere maintenance [26,27]. It remains however possible that the p53-dependent repression of additional genes, such as *Dclre1b*, mutated in dyskeratosis congenita, or *Fancd2* [65], might also affect telomere maintenance. Likewise, increased p53 activity was reported to partially phenocopy Diamond-Blackfan anemia (DBA), through mechanisms that remained unknown [28]. Our finding that *Tsr2*, a gene mutated in DBA [66], is repressed by p53 and DREAM provides a possible explanation for DBA-like phenotypes consecutive to germline p53 activation, but the p53-dependent repression of *Fanca* might also contribute to altered ribosome biogenesis [41]. Altogether, these data suggest that an increased p53 activity may cause bone marrow failure through several possible mechanisms, by promoting the DREAM-mediated repression of many genes. Although this complexity may hamper the identification of the most clinically relevant p53-DREAM targets, it might also account for the partial phenotypic overlap between bone marrow failure syndromes of distinct molecular origins, as discussed previously [25]. Indeed, defects in telomere maintenance, DNA repair or ribosome function would all lead to p53 activation [67–69], and the subsequent DREAM-mediated gene repression might have similar downstream consequences, leading to common clinical traits. Furthermore, our analyses indicated that many targets of the p53-DREAM pathway are associated with microcephaly or cerebellar hypoplasia, also suggesting that a DREAM-mediated concomitant downregulation of multiple genes might contribute to these pathological processes. Consistent with this possibility, the Zika virus (ZIKV) is known to cause p53 activation in cortical neural progenitors and microcephaly [49,50], and genetic analyses in ZIKV-infected mice indicated that variations in clinical severity and brain pathology between different mouse strains were driven by multiple host genes with small effects [70].

Our analysis suggests that many targets of the p53-DREAM pathway are associated with syndromes of abnormal hematopoiesis or brain development. To get a more precise evaluation of this association, we searched for genetic disorders that might be caused by the mutation of any the 151 candidate p53-DREAM targets for which putative DBS were identified. According to OMIM, the online catalog of human genes and genetic disorders (https://www.omim.org), 106/151 genes were mutated in a hematological or neurological disorder. Among these, 25 were mutated in syndromes characterized by anemia, lymphopenia, neutropenia or thrombocytopenia and 77 in syndromes with microcephaly, cerebellar hypoplasia or hypoplasia of cerebellar vermis, including 13 associated with both types of symptoms (Tables 3 and S36). Among these 13 genes is notably *Nuf2*, whose mutations were initially shown to cause microcephaly [71] but were later also associated with bone marrow failure [72]. Furthermore, out of 317 potential DREAM targets, 58 were associated with both blood- and brain-related gene ontology terms (Tables S17, S26). This suggests that it might be worthwhile to systematically search for hematopoietic anomalies in patients with syndromes of abnormal brain development and, conversely, to check for neurological anomalies in patients with syndromes of abnormal hematopoiesis.

**Table 3.**
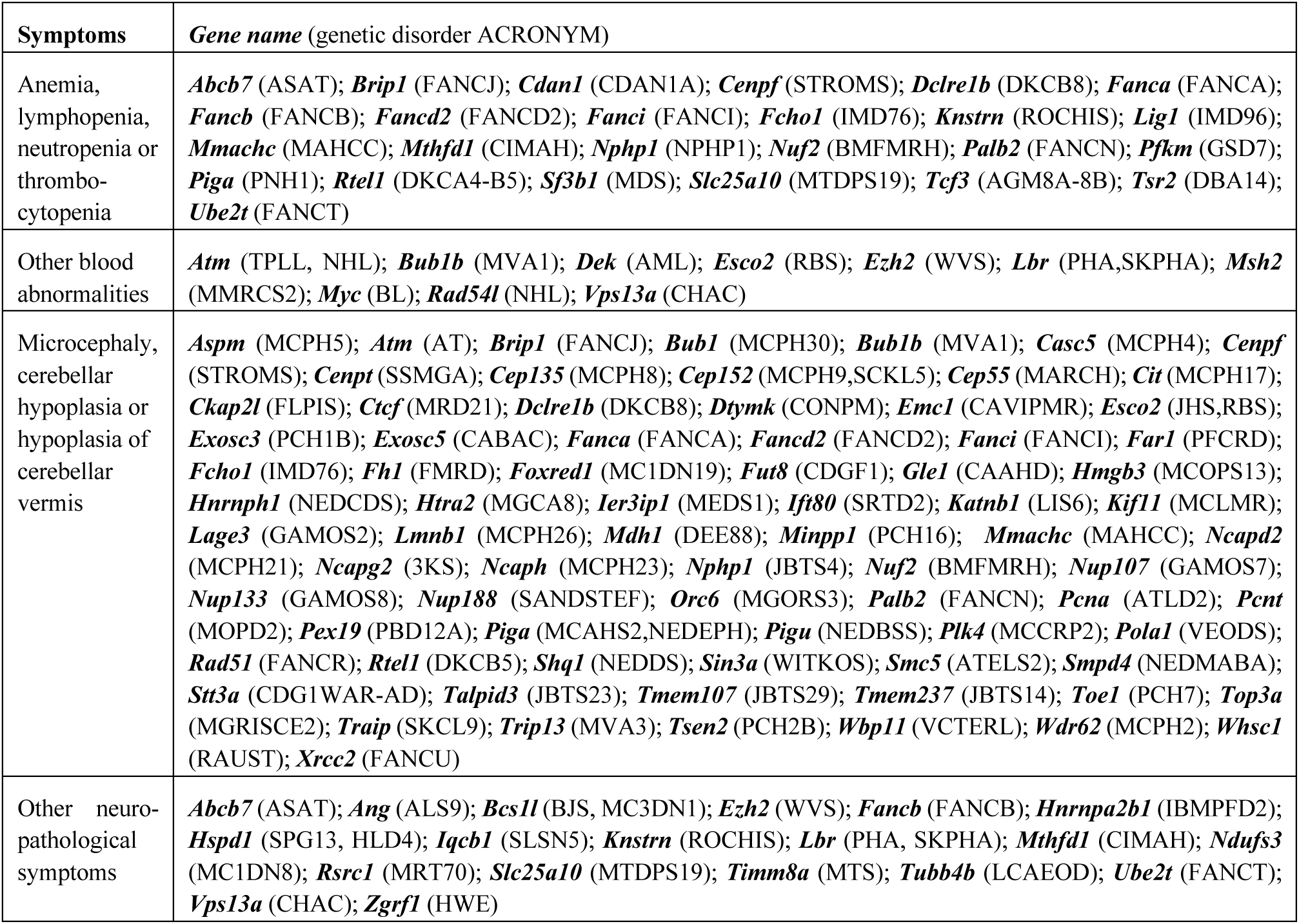
Candidate p53-DREAM targets mutated in genetic disorders with hemato- or neuro-pathological symptoms. Acronyms for genetic disorders are 3KS: Khan-Khan-Katsanis syndrome; AGM: Agammaglobulinemia; AML: acute myeloid leukemia; ALS: amyotrophic lateral sclerosis; ASAT: anemia sideroblastic and spinocerebellar ataxia; AT: ataxia telangectasia; ATELS: Atelis syndrome; ATLD: AT-like disorder; BJS: Bjornstad syndrome; BL: Burkitt lymphoma; BMFMRH: Bone marrow failure with microcephaly and renal hypoplasia; CAAHD: congenital arthrogryposis with anterior horn cell disease; CABAC: cerebellar ataxia, brain abnormalities and cardiac conduction defects; CAVIPMR: cerebellar atrophy, visual impairment and psychomotor retardation; CDAN: congenital dyserythropoietic anemia; CDG: congenital disorder of glycosylation; CDGF: CDG with defective fucosylation; CHAC: chorea-acanthocytosis; CIMAH: combined immunodeficiency and megaloblastic anemia with or without hyperhomocysteinemia; CONPM: childhood-onset neurodegeneration with progressive microcephaly; DBA: Diamond-Blackfan anemia; DEE: developmental and epileptic encephalopathy; DKC: dyskeratosis congenita; FANC: Fanconi anemia; FLPIS: Filippi syndrome; FMRD: fumarase deficiency; GAMOS: Galoway-Mowat syndrome; GSD: glycogen storage disease; HLD: hypomyelinating leukodystrophy; HWE: hot water epilepsy; IBMPFD: inclusion body myopathy with early-onset Paget disease with or without fronto-temporal dementia; IMD: immunodeficiency; JBTS: Joubert syndrome; JHS: Juberg-Hayward syndrome; LCAEOD: Leber congenital amaurosis with early-onset deafness; LIS: lissencephaly; MAHC: methylmalonic aciduria and homocystinuria; MARCH: multinucleated neurons, anhydramnios, renal dysplasia, cerebellar hypoplasia and hydranencephaly; MC[1,3]DN: mitochondrial complex [I or III] deficiency, nuclear type; MCAHS: multiple congenital anomalies-hypotonia-seizures syndrome; MCCRP: microcephaly and chorioretinopathy; MCLMR: microcephaly, with or without chorioretinopathy, lymphedema or mental retardation; MCOPS: syndromic microphthalmia; MCPH: primary microcephaly; MDS: myelodysplastic syndrome; MEDS: microcephaly, epilepsy and diabetes syndrome; MGCA: 3-methylglutaconic aciduria; MGORS: Meier-Gorlin syndrome; MGRISCE: microcephaly, growth restriction and increased sister-chromatid exchange; MMRCS: mismatch repair cancer syndrome; MOPD: microcephalic osteodysplastic primordial dwarfism; MRD or MRT: intellectual developmental disorder; MTDPS: mitochondrial DNA depletion syndrome; MTS: Mohr-Tranebjaerg syndrome; MVA: mosaic variegated aneuploidy syndrome; NED: neurodevelopmental disorder; NEDBSS: NED with brain anomalies, seizures and scoliosis; NEDCDS: NED with craniofacial dysmorphism and skeletal defects; NEDDS: NED with dystonia and seizures; NEDEPH: NED with epilepsy and hemochromatosis; NEDMABA: NED with microcephaly, arthrogryposis, and structural brain anomalies; NHL: non-Hodgkin lymphoma; NPHP: nephronophthisis; PBD: peroxisome biogenesis disorder; PCH: pontocerebellar hypoplasia; PFCRD: peroxisomal fatty acyl-CoA reductase 1 disorder; PHA: Pelger-Huet anomaly; PNH: paroxysmal nocturnal hemoglobinuria; RAUST: Rauch-Steindl syndrome; RBS: Robert-Sc phocomelia syndrome; REYNS: Reynolds syndrome;ROCHIS: Roifman-Chitayat syndrome; SANDSTEF: Sandestig-Stefanova syndrome; SCKL: Seckel syndrome; SKPHA: rhizomelic skeletal dysplasia with or without PHA; SLSN: Senior-Loken syndrome; SPG: spastic paraplegia; SRTD: short-rib thoracic dysplasia; SSMGA: short stature, microcephaly with genital anomalies; STROMS: Stromme syndrome; TPLL: T-cell prolymphocytic leukemia; VCTERL: vertebral, cardiac, tracheoesophageal, renal, and limb defects; VEODS: Van Esch-O’Driscoll syndrome; WITKOS: Witteveen-Kolk syndrome; WVS: Weaver syndrome. See Table S36 for additional details.

The p53-DREAM targets we identified are likely to be overexpressed in cells with mutant p53, a frequent alteration in cancer cells. For some p53-DREAM targets, such an overexpression may promote tumorigenesis. For example, TRIP13 was shown to promote cancer cell proliferation and epithelial-mesenchymal transition (EMT) in various tumor types [73–78] and we identified here a functionally relevant DBS regulating *Trip13* expression. In cells with mutant p53, an increase in TRIP13 expression might thus be one of the mechanisms favoring EMT. Importantly, many of the p53-DREAM targets we identified play a role in brain development, suggesting that the impact of a loss or attenuation of the p53-DREAM pathway might be particularly relevant for brain tumorigenesis. In support of this possibility, the chromatin regulator bromodomain-containing protein 8 (BRD8) was recently shown to attenuate p53 in glioblastoma [79] and we observed, in glioblastoma cells with high BRD8 levels [80], an overall increased expression for the 77 p53-DREAM targets associated with microcephaly or cerebellar hypoplasia (Figure 4a, Table S37). Furthermore *CENPF*, *ASPM* and *CASC5* are known to contribute to phenotypic variation in glioblastoma neoplastic cells [81], and they were among the 8 genes most affected by BRD8 levels (Figure 4b, Table S37).

**Figure 4.**
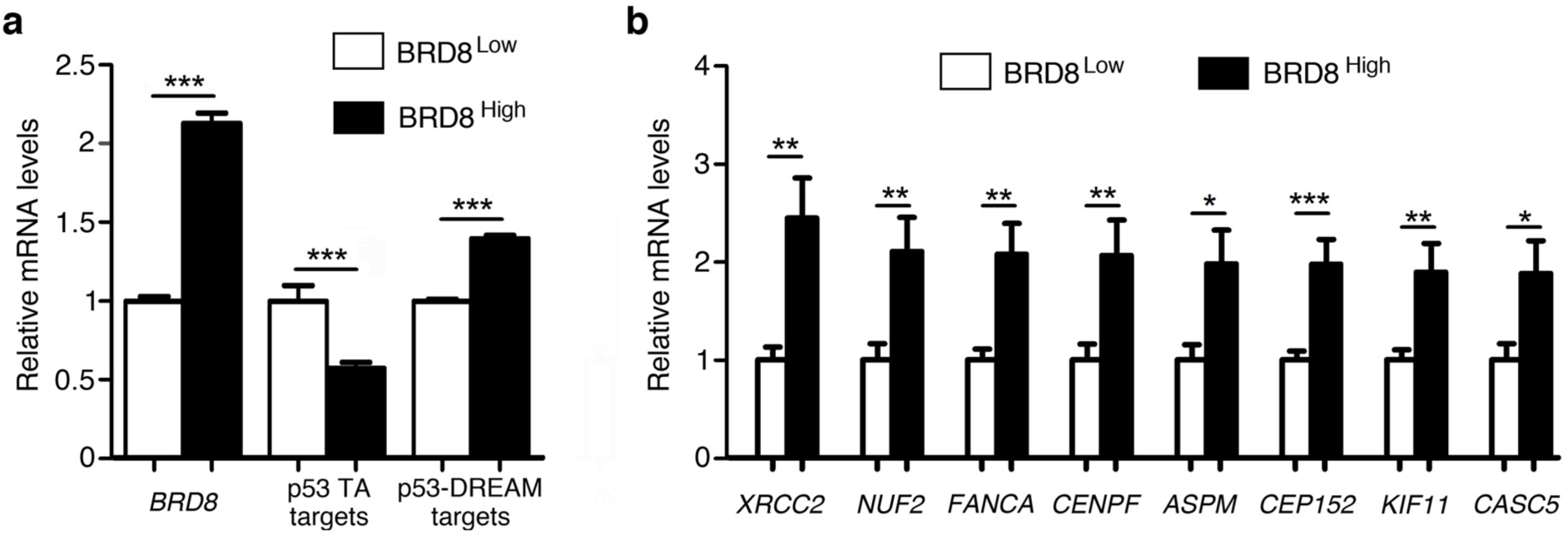
In glioblastoma cells, high BRD8 levels correlate with an increased expression of brain-related p53-DREAM targets. **(a)** Glioblastoma with high BRD8 levels exhibit an overall increased expression of brain-related p53-DREAM target genes. RNAseq data from glioblastoma cells isolated from patient specimens (GSE121720) were stratified according to BRD8 mRNA levels as previously described [79], and for each gene the average expression levels in tumors expressing low BRD8 levels was assigned a value of 1. The relative expression for *BRD8*, five p53-transactivated (TA) target genes (*CDKN1A*, *MDM2*, *BAX*, *GADD45A*, *PLK3*) and the 77 p53-DREAM targets associated with microcephaly or cerebellar hypoplasia (from Table 3) are shown. Tumors with high BRD8 expression levels exhibit decreased expression levels of p53 transactivated targets and increased levels of p53-DREAM targets. Data are from 23 samples per group for *BRD8*, 115 values per group (23 samples x 5 genes) for p53 transactivated targets and 1,771 values per group (23 samples x 77 genes) for p53-DREAM targets. **(b)** *CENPF*, *ASPM* and *CASC5* are among the 8 p53-DREAM target genes whose expression is most affected by BRD8 levels in glioblastoma cells. Data for the indicated genes (with 23 samples per group) were retrieved from dataset GSE121720 and analyzed as in (a). \*\*\**P<*0.001, \*\**P<*0.01, \**P<*0.05 in Student’s *t* tests. See Table S37 for additional details.

Altogether, this analysis expands our knowledge of the p53-DREAM pathway and notably indicates that this pathway regulates many genes implicated in bone marrow failure syndromes, neurodevelopmental disorders and cancer, suggesting an explanation for the variety of clinical symptoms that might result from its deregulation. Furthermore, our positional frequency matrices, which were useful to identify functionally relevant DREAM binding sites in genes associated with blood- or brain-related syndromes, should be considered to analyze the promoters of additional DREAM targets, implicated in other pathologies.

## Materials and Methods

### Transcriptome data comparisons

We analyzed the gene expression data from Hoxa9-ER expressing hematopoietic stem and progenitor cells grown in the presence of tamoxifen, or in differentiated cells 5 days after tamoxifen withdrawal, a microarray study relying on 45,101 probes corresponding to 20,627 genes (Gene Expression Omnibus #GSE21299) [34]. For each probe, we calculated the inverse of Log2 from robust multi-average values. The obtained average (from triplicates) for cells with tamoxifen was given a value of 1, and the ratios before and after tamoxifen withdrawal were calculated. For each gene, we took the probe leading to the highest repression ratio into account and selected those downregulated at least 1.5-fold upon tamoxifen withdrawal. Among those genes, we identified targets downregulated by human and/or mouse p53 by consulting p53 regulation scores in the Target gene regulation database (www.targetgenereg.org) [23]. Relative expression data were graphed with Microsoft Excel, by using a two-color scale and conditional coloring.

### Gene ontology analyses

To identify genes associated with bone marrow failure, we first used the Gene Ontology enRIchment anaLysis and visuaLizAtion tool (GOrilla, Technion) [35]. Out of 20,627 genes analyzed by microarray, 17,461 were associated with a Gene Ontology term according to GOrilla. A downregulation of at least 1.5-fold upon tamoxifen withdrawal was observed for 6,880 probes corresponding to 4,571 genes, of which 3,631 were associated with a GO term. Enrichment analyses were carried out by comparing the unranked list of genes downregulated at least 1.5 fold (target) to the full list of genes (background), with ontology searches for biological processes or molecular function and default P value settings (10^-3^). Independently, for both blood- and brain-related genes, we used the GO lists from the Human Phenotype Ontology website of the Jackson Laboratory [43].

### ChIP-seq data analyses

We used the peak browser from ChIP-Atlas (https://chip-atlas.org/peak_browser) [38] to search for E2f4 and Lin9 binding on the Mus musculus (mm10) genome, or E2F4 and LIN9 binding on the Homo sapiens (hg38) genome, and visualized results on the Integrative genomics viewer (IGV_2.12.2) [82]. Peaks from all cell types were analyzed, and those with the highest binding score and minimal distance from the transcription start site were selected for. ChIP binding scores were graphed with Microsoft Excel, by using a two-color scale and conditional coloring.

### RNAseq data analyses

To screen for the most relevant candidate p53-DREAM targets, we analyzed publicly available datasets GSE171697 and GSE204924 for blood ontology-related genes, and datasets GSE78711 and GSE80434 for brain ontology-related genes. In addition, we analyzed the dataset GSE121720, containing RNAseq data from glioblastoma cells isolated from patient specimens. This dataset contains 92 samples, which were ranked according to BRD8 expression levels, and the top and bottom 25% samples were assigned as the BRD8^high^ and BRD8^low^ groups, as previously described [79,80]. Data for the 77 p53-DREAM targets associated with microcephaly or cerebellar hypoplasia (from Table 3) were then retrieved from the dataset and analyzed.

### Search for putative DREAM binding sites

To search for putative DREAM binding sites, we used PWMScan (https://ccg.epfl.ch/pwmtools/pwmscan.php [51]) with a custom positional frequency matrix from 10, 21 or 22 murine functional DBS (for details see Figure S7), on both the mouse (mm10) and human (hg38) genomes, with a P value threshold of 10^-3^ for cut-off. The putative DREAM binding sites identified were then analyzed for sequence conservation between mouse and human genomes and classified according to PFM score and number of mismatches between the two species at positions 2-6 (for CDE or E2F) and 11-16 (for CHR or CLE) of the DREAM binding sites. For 151 genes, the identified putative DBS, as well as putative E2F or CHR sites reported in the Transcriptional regulation database, were mapped relative to ChIP peaks (or transcriptional start sites) by using the Integrative Genomics Viewer (igv version 2.12.2).

### Cells and cell culture reagents

NIH3T3 cells, or mouse embryonic fibroblasts (MEFs) isolated from 13.5 days post-coitum embryos and cultured for <5 passages, were cultured in a 5% CO_2_ and 3% O_2_ incubator, in DMEM Glutamax (GIBCO), with 15% FBS (PAN Biotech), 100 μM 2-mercaptoethanol (Millipore), 0.01 mM Non-Essential Amino-Acids and Penicillin/Streptomycin (GIBCO). Cells were treated for 24h with 10 μM Nutlin 3a (Sigma-Aldrich).

### Quantitative RT-PCR

Total RNAs were extracted using nucleospin RNA II (Macherey-Nagel), reverse-transcribed using superscript IV (Invitrogen), and real-time quantitative PCRs were performed on an ABI PRISM 7500 using Power SYBR Green (Applied Biosystems) as previously described [26]. Primer sequences are listed in Table S38.

### Luciferase assays

For each tested gene, a 1-1.5 kb fragment of the promoter containing at its center the putative DBS was cloned upstream a luciferase reporter gene in the backbone of a PGL3 basic vector (Promega). For all tested DBS, to prevent DREAM binding we used PCR mutagenesis and mutated the putative binding site into the following sequence: 5’-AAATAA(NNNN)AGACTG-3’, with (NNNN) corresponding to DNA spacer sequences that were not mutated. We used lipofectamine 2000 to transfect 10^6^ NIH-3T3 MEFs with 3 μg of the luciferase plasmid with a WT DBS or its mutant counterpart and 30 ng of a renilla luciferase expression plasmid (pGL4.73, Promega) for normalization, and treated or not with 10 μM Nutlin 3a. Transfected cells were incubated for 24 h then trypsinized, resuspended in 75 μl culture medium with 7.5% FBS and transferred into a well of an optical 96-well plate (Nunc). The dual-glo luciferase assay system (Promega) was used according to the manufacturer’s protocol to lyse the cells and read firefly and renilla luciferase signals. Results were normalized, then the average luciferase activity in untreated cells transfected with a WT Promoter were assigned a value of 1.

### Statistical Analyses

Student’s *t* tests were used to analyze differences between undifferentiated and differentiated bone marrow cells, between WT and p53^Δ31/Δ31^ bone marrow cells, untreated and Nutlin-treated cells, between WT or mutant promoters in luciferase assays, and between glioblastoma with low or high BRD8 levels. Analyses were performed using Graphpad Prism 5, and values of P<0.05 were considered significant. Hypergeometric testing of DREAM binding site distributions was performed with the Keisan calculator (keisan.casio.com).

## Acknowledgements

F.T. received funding form the Fondation ARC (projet) and the Ligue Nationale Contre le Cancer (Comité Ile-de-France); J.R. and E.E. are PhD fellows of the Ministère de la Recherche. We thank L. Chen for technical assistance, J. Josephides for suggestions in data analysis, and A. Fajac and B. Bardot for critical reading of the manuscript. Author contributions: conceptualization and project supervision: F.T.; investigation: J.R., V.L., F.T., C.D., E.E.; formal analysis: H.E.; funding acquisition: F.T.; manuscript preparation: F.T., J.R., V.L. The authors declare that they have no conflict of interest.

