## Supplemental Figures for "A systematic approach identifies p53-DREAM target genes associated with blood or brain abnormalities"

**Supplementary Figures.**

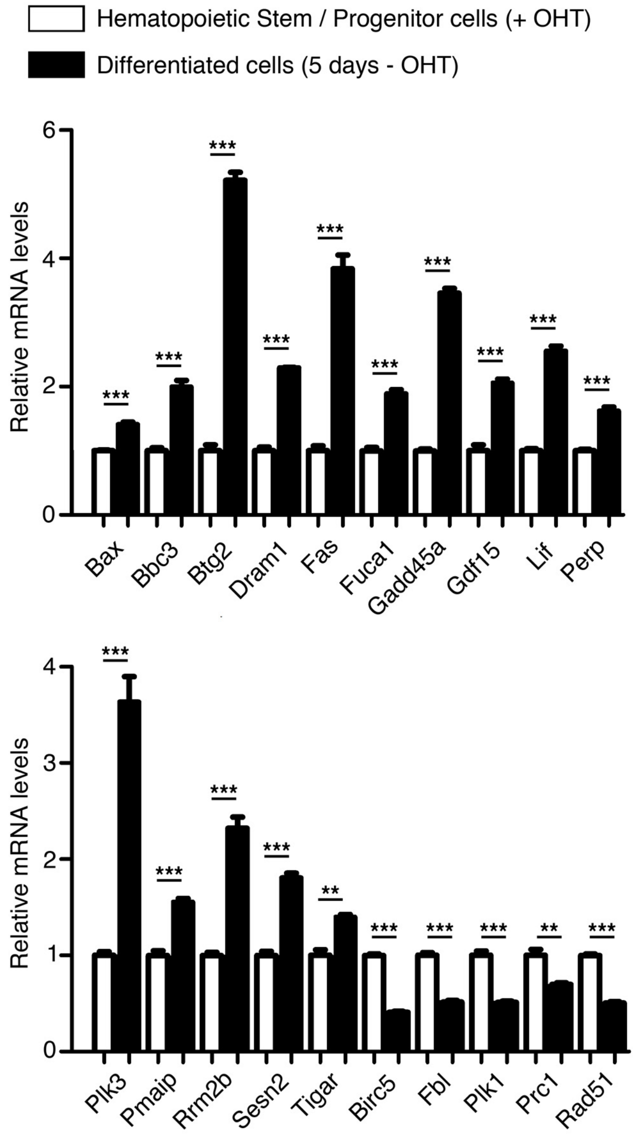


**Figure S1. Additional evidence that Hoxa9-ER expressing bone marrow cell differentiation correlates with p53 activation.**

Robust-multi average values for expression of the indicated genes were extracted from transcriptome data of Hoxa9-ER expressing hematopoietic stem and progenitor cells (grown in the presence of tamoxifen) or differentiated cells (5 days after tamoxifen withdrawal). Average values (from triplicates) in cells with tamoxifen were given a value of 1. Upon p53 activation, *Bax*, *Bbc3/Puma*, *Btg2*, *Dram1*, *Fas*, *Fuca1*, *Gadd45a*, *Gdf15*, *Lif*, *Perp*, *Plk3*, *Pmaip*, *Rrm2b*, *Sesn2* and *Tigar* are known to be transactivated, whereas *Birc5/Survivin*, *Fbl*, *Plk1*, *Prc1* and *Rad51* are known to be downregulated. Means + s.e.m. are shown; ***P<0.001, **P<0.01 by Student's t test.

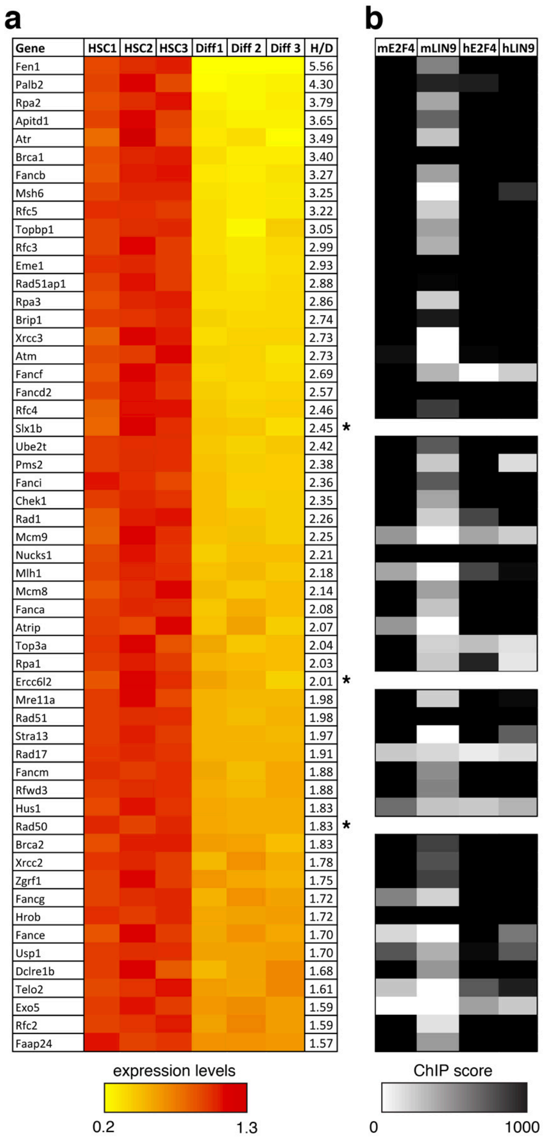


**Figure S2. Fanconi-related genes downregulated upon bone marrow cell differentiation, and their potential regulation by DREAM.**

(**a**) Expression values, represented as described in Figure 1b, for 55 genes related to the Fanconi anemia DNA repair pathway, of which 52 are regulated by p53. (**b**) ChIP scores of E2F4 or LIN9 binding in mouse (m) or human (h) cells for the 52 Fanconi-related, p53-regulated genes, represented as in Figure 1c.

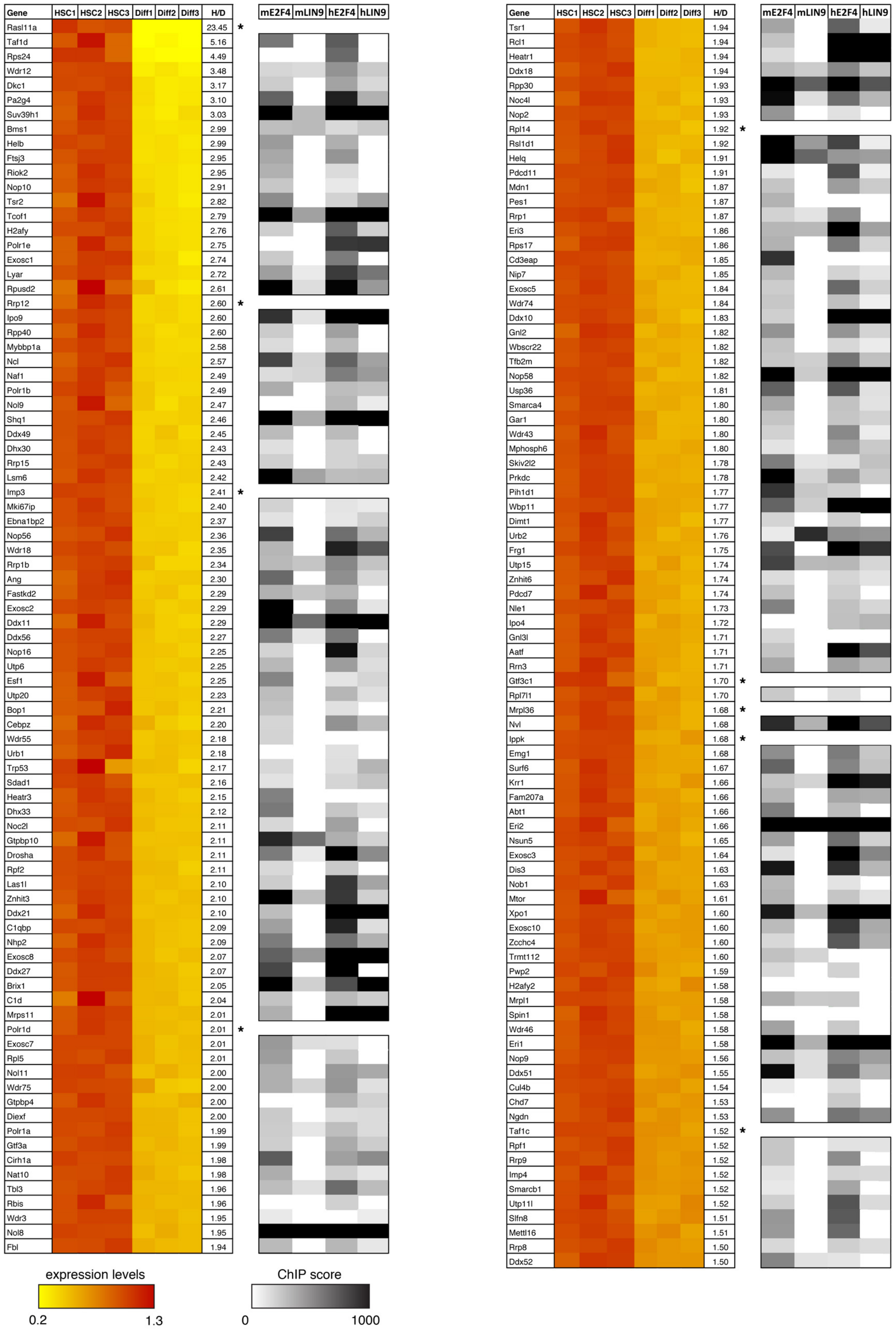


**Figure S3. Ribosome-related genes downregulated upon bone marrow cell differentiation, and their potential regulation by DREAM.**
Expression values and highest E2F4 and LIN9 ChIP binding scores for 171 ribosome-related genes, represented as described in Figure 1b-c.

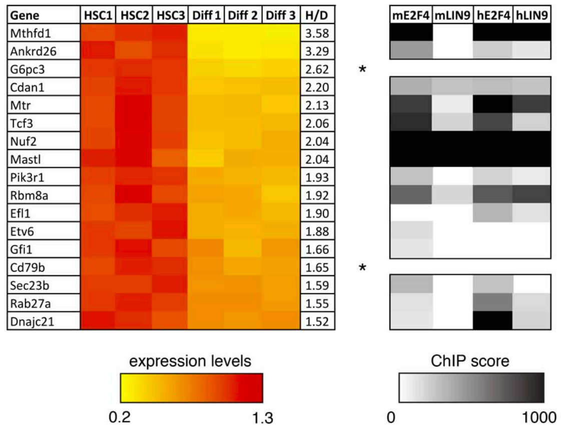


**Figure S4. Other genes associated with bone marrow failure syndromes, downregulated upon bone marrow cell differentiation, and their potential regulation by DREAM.** Expression values and highest E2F4 and LIN9 ChIP binding scores for 17 genes mutated in other bone marrow failure syndromes, represented as described in Figure 1b-c.

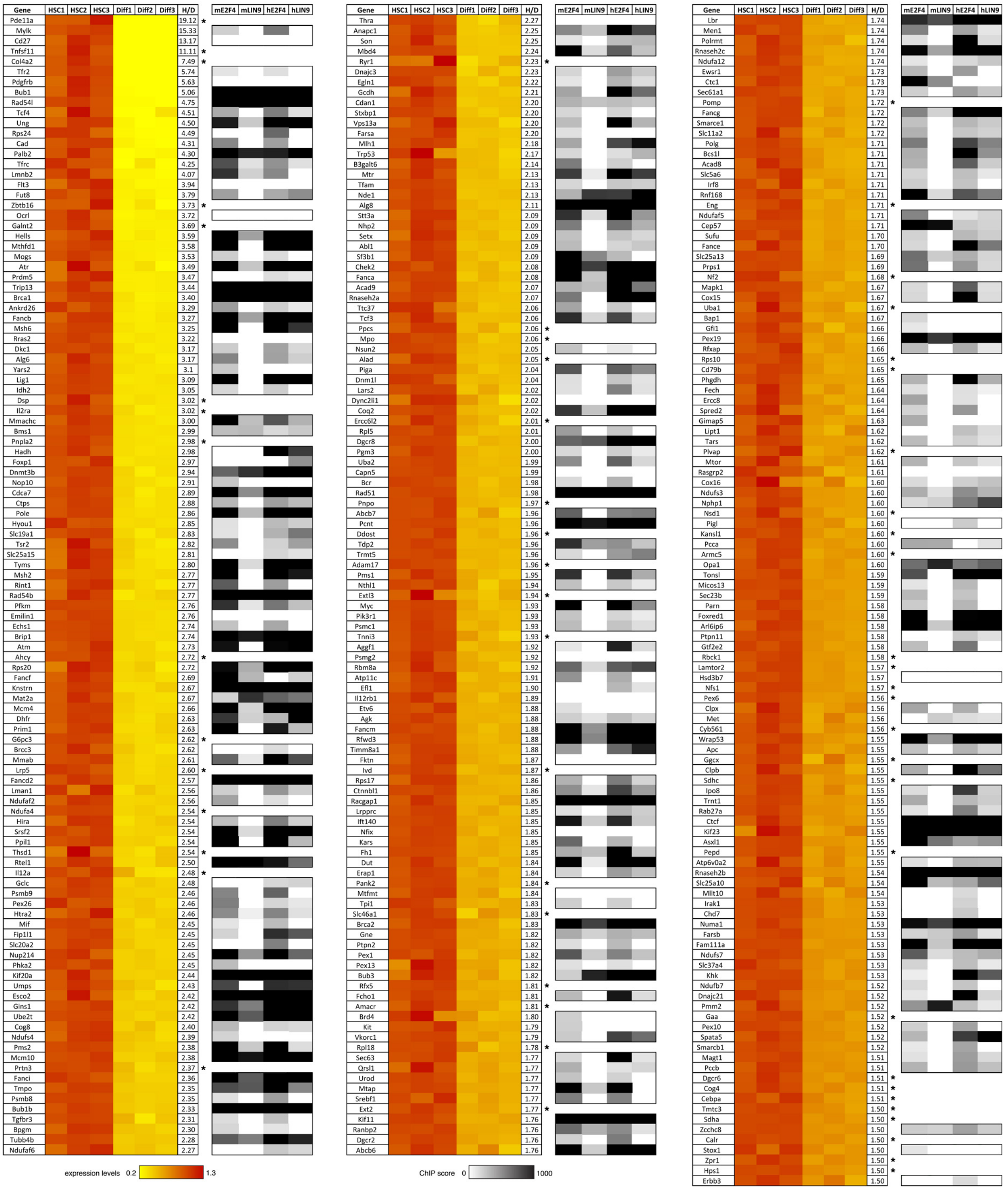


**Figure S5. Genes associated with abnormalities in blood or blood-forming tissues and downregulated upon bone marrow cell differentiation, and their potential regulation by DREAM.**

Expression values and highest E2F4 and LIN9 ChIP binding scores for 336 genes associated with abnormalities in blood and blood-forming tissues (according to the Human Phenotype Ontology website of the Jackson laboratory, ontology term #HP:001871), represented as described in Figure 1b-c.

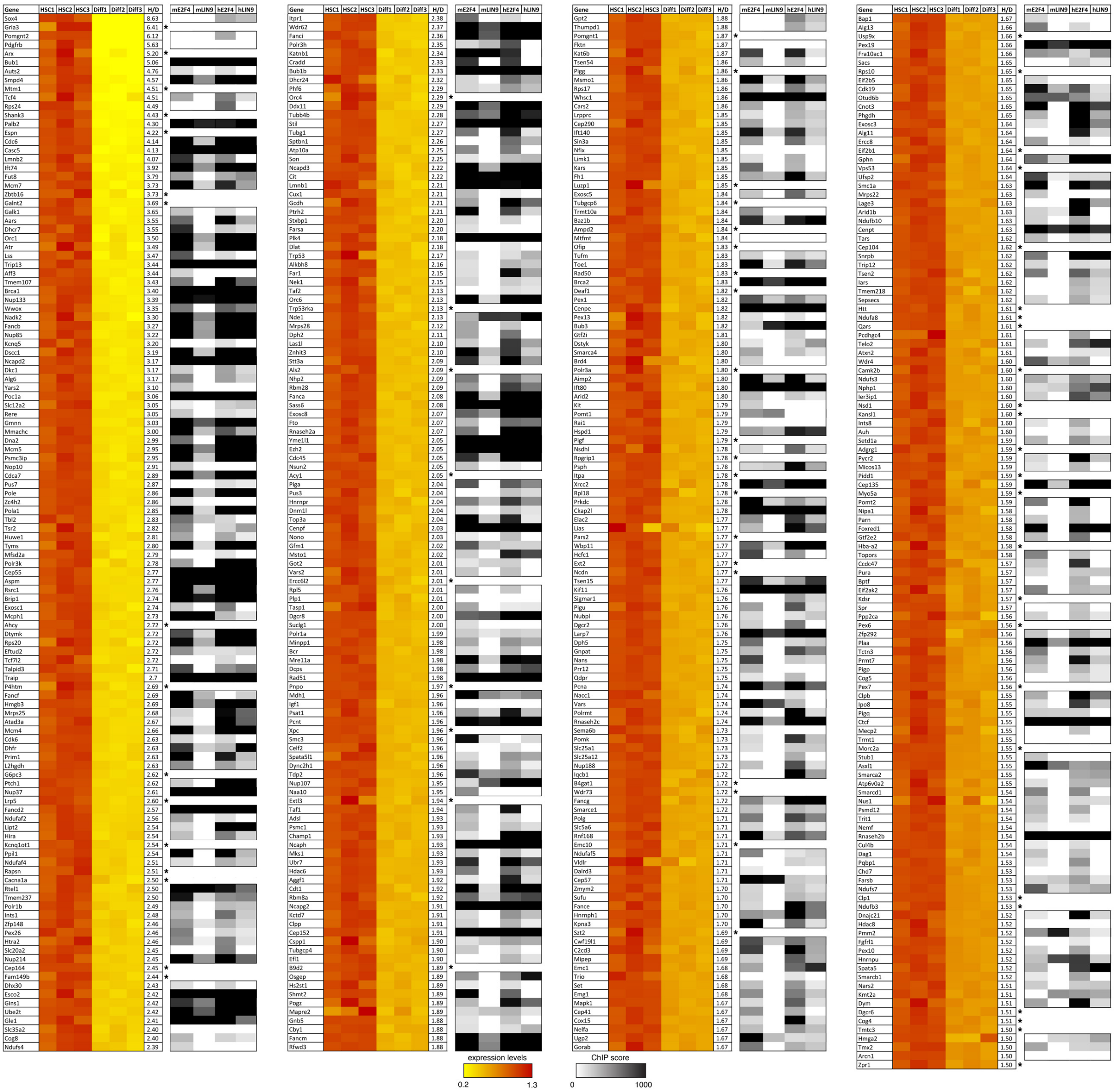


**Figure S6. Genes associated with microcephaly or cerebellar hypoplasia and downregulated upon bone marrow cell differentiation, and their potential regulation by DREAM.**

Expression values and highest E2F4 and LIN9 ChIP binding scores for 474 genes associated with microcephaly or cerebellar hypoplasia (according to the Human Phenotype Ontology website of the Jackson laboratory, ontology terms #HP0000252 and HP:0007360), represented as described in Figure 1b-c.

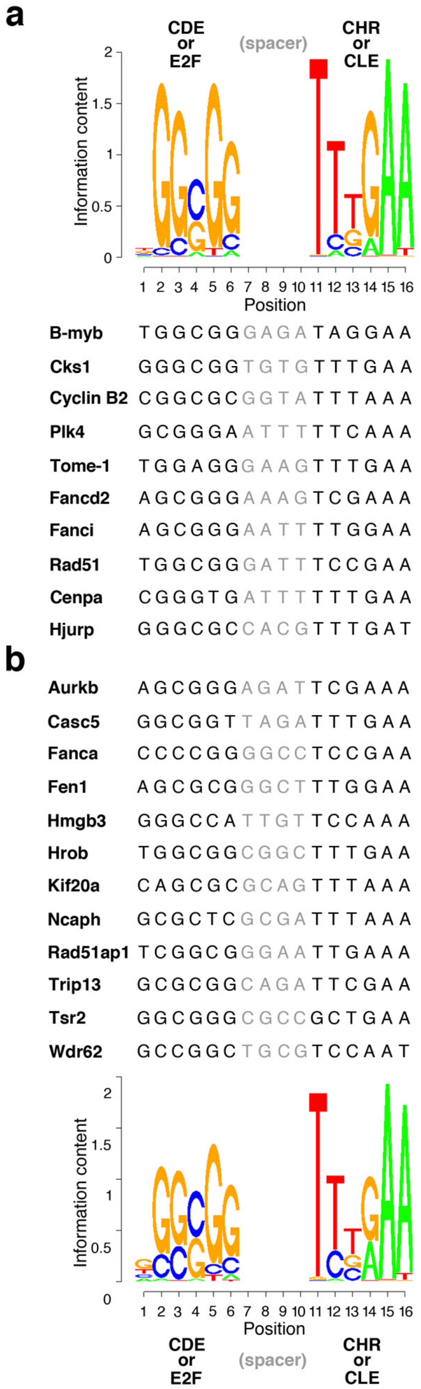


**Figure S7. Positional frequency matrices used to search for DREAM binding sites.**
**(a)** PFM10, a positional frequency matrix from 10 experimentally validated DREAM binding sites (DBS). The logo for PFM10 (top) and DNA sequences from the 10 experimentally validated murine DBS used to generate it (bottom) are shown. Spacer DNA sequences between GC-rich (CDE or E2F) and AT-rich (CHR or CLE) elements were not used to define the matrix. **(b)** PFM22, a positional frequency matrix from 22 experimentally tested DBS. The DNA sequences of 12 additional murine DBS tested in this study (top) were added to the first 10 DBS to define PFM22 (bottom). Spacer DNA sequences between GC-rich (CDE or E2F) and AT-rich (CHR or CLE) elements were not taken into account to define the matrix. Compared to PFM10, PFM22 notably introduces minor nucleotides at positions 2 (A), 5 (C), 6 (T) and 11 (G), while decreasing the frequency of rare nucleotides at positions 4 (A) and 12 (A).

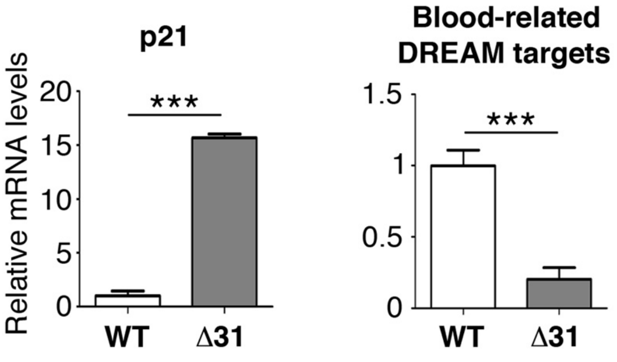


**Figure S8. Bone marrow cells from p53$^{Δ31/Δ31}$ mice exhibit increased p21 gene expression, and a decreased expression of blood-related candidate p53-DREAM targets.**
mRNAs extracted from the bone marrow cells of wild-type (WT) and p53$^{Δ31/Δ31}$ (Δ31) mice were quantified using real-time PCR, normalized to control mRNAs, then the amount in WT untreated cells was assigned a value of 1. The blood–related p53-DREAM target genes tested are the 8 genes reported in Figure 2a-c : *Aurkb*, *Fanca*, *Fen1*, *Hrob*, *Kif20a*, *Rad51ap1*, *Trip13* and *Tsr2*. Results from 2-3 mice per genotype, thus 16-24 values per group of DREAM target genes. ***P<0.001 by Student's t test.

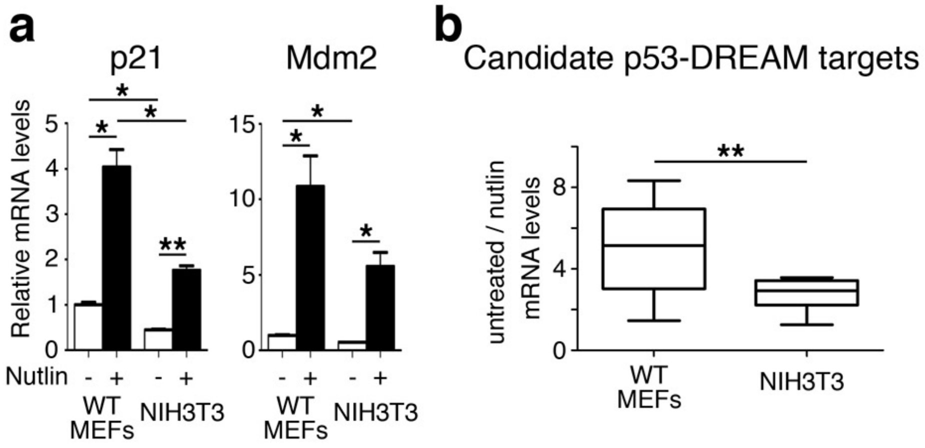


**Figure S9. The p53-DREAM pathway is attenuated in NIH3T3 cells.**
**(a)** The transactivation of p53 target genes is decreased in NIH3T3 cells. mRNAs from primary wild-type (WT) MEFs or NIH3T3 cells, untreated or treated with 10 mM Nutlin for 24h, were quantified using real-time PCR, normalized to control mRNAs, then the amount in WT untreated MEFs was assigned a value of 1. Means + s.e.m. from 2-3 independent experiments are shown. **(b)** Upon p53 activation, the repression of DREAM targets is less pronounced in NIH3T3 cells. The mRNAs for 12 candidate p53-DREAM targets (*Aurkb*, *Casc5*, *Fanca*, *Fen1*, *Hmgb3*, *Hrob*, *Kif20a*, *Ncaph*, *Rad51ap1*, *Trip13*, *Tsr2*, *Wdr62*), extracted from primary wild-type (WT) MEFs or NIH3T3 cells, untreated or treated with 10 mM Nutlin for 24h, were quantified using real-time PCR and normalized to control mRNAs as above, and ratios of nutlin-induced repression were calculated. For each tested gene, results are from 2-3 independent experiments. **P<0.01, *P<0.051 by Student's t test.

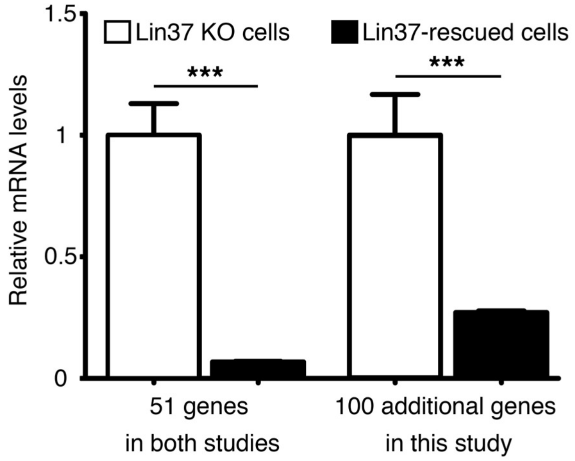


**Figure S10. The expression of candidate p53-DREAM targets is decreased in Lin37-rescued cells.**

For the 151 genes with putative DREAM binding sites (see Table 2), we extracted RNAseq data from dataset GSE97716, in LIN37 KO cells and LIN37-rescued cells. Data were from 2 different KO clones and 2 rescued clones, with 2-3 values per clone for each gene. For each gene, average expression values were calculated and a value of 1 was attributed for expression levels in LIN37 KO cells. Out of 151 genes, only 51 genes were previously reported to be DREAM targets based on their differential expression upon Lin37 reintroduction (for details, see main text and Table S35). Nevertheless, the other 100 genes were also downregulated upon Lin37 reintroduction, albeit with a smaller fold decrease. ***P<0.001 by Student's t test.

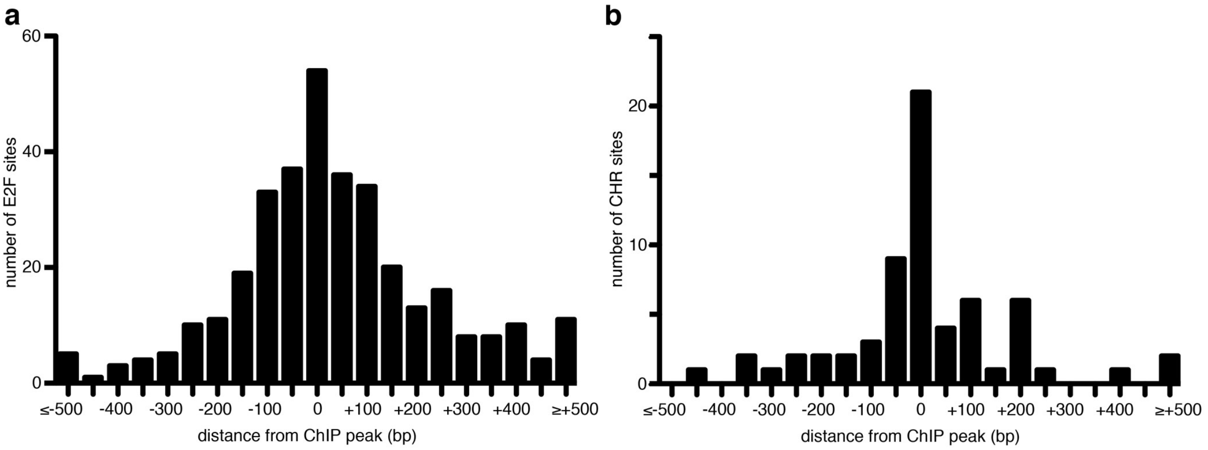


**Figure S11. Mapping of E2F and CHR motifs from the TGR database relative to ChIP peaks for DREAM subunits.**

**(a)** The 342 E2Fs motifs reported in the TGR database for the 151 genes listed in Table 2 were mapped relative to the ChIP peaks of E2F4 and/or LIN9 binding, in 50 bp windows. **(b)** The 64 CHR motifs reported in the TGR database for the 151 genes listed in Table 2 were mapped relative to the ChIP peaks of E2F4 and/or LIN9 binding, in 50 bp windows.
